# Detecting Extrachromosomal DNA from Routine Histopathology

**DOI:** 10.64898/2026.02.27.708546

**Authors:** Muhammad Anwaar Khalid, Michael Gratius, Christopher Brown, Raneen Younis, Zahra Ahmadi, Lukas Chavez

## Abstract

Extrachromosomal DNA (ecDNA) is a major driver of oncogene amplification, tumour heterogeneity and poor clinical outcomes [1–3], yet its detection relies on specialised genomic assays that are not integrated into routine diagnostics. Here, we show that ecDNA status can be inferred directly from standard haematoxylin and eosin–stained whole-slide pathology images. We develop an end-to-end, weakly supervised deep learning framework that aggregates thousands of high-magnification patches per slide with slide-level augmentation and interpretable attention. Across twelve cancer types from The Cancer Genome Atlas, the approach identifies tumours with genomic amplifications and, critically, distinguishes ecDNA-amplified from chromosomally amplified or non-amplified tumours, with the strongest signal in glioblastoma. Attention maps localise regions enriched for nuclei with altered chromatin intensity and texture, and predicted ecDNA status recapitulates its adverse association with survival. These results indicate that ecDNA amplifications leave reproducible histomorphologic footprints detectable by routine pathology, enabling scalable screening to prioritise tumours for confirmatory molecular testing.

## Main

Extrachromosomal DNA (ecDNA) comprises circular, acentric DNA elements that segregate unevenly through cell division, can achieve high oncogene copy number, and remodel transcriptional programs, thereby intensifying intra-tumour heterogeneity and resistance to therapy [2–4]. Across adult and paediatric cancers, ecDNA associates with aggressive phenotypes and inferior survival relative to chromosomal amplification or the absence of focal amplification [5–8]. Recent evidence suggests that ecDNA formation is not merely a late-stage adaptation but can arise early during malignant transformation, marking the transition from high-grade dysplasia to invasive disease [9, 10]. These observations position ecDNA as both a fundamental driver of cancer biology and a potentially actionable biomarker for risk stratification and therapeutic decision-making [11]. Yet ecDNA assessment is uncommon in clinical practice because reliable detection requires karyotyping or fluorescence in situ hybridisation for selected loci, or whole-genome sequencing with specialised reconstruction [12], modalities that are costly, time-consuming, and not routinely available at diagnosis. This gap between biological importance and clinical accessibility limits population-scale studies and timely risk stratification for patients whose tumours may be ecDNA-driven. Routine hematoxylin and eosin (H&E) whole-slide images (WSIs), by contrast, are acquired for nearly all solid tumours and capture rich morphology that reflects underlying genetic and microenvironmental states [13, 14]. In pancreatic ductal adenocarcinoma, *MYC* amplification on ecDNA drives reversible shifts from glandular to solid and cribriform growth patterns, with *MYC* -ecDNA–rich regions co-localising with distinct histological architectures [15]. In glioblastoma, ecDNA-positive tumours are enriched for mesenchymal-like malignant states and hypoxia-associated vascular niches, biological programs that localise to canonical histological features [16]. These findings support the hypothesis that ecDNA biases tumours toward specific, spatially organised morphological states, raising the possibility that ecDNA status may be detectable from routine H&E histopathology.

## A multi-cancer cohort with matched histology and amplification labels

We assembled a multi-cancer patient cohort by integrating histopathology slides with publicly available genomic amplification annotations (Figure 1a). Digitized H&E–stained WSIs were retrieved from The Cancer Genome Atlas (TCGA) [17] via the Genomic Data Commons (GDC) Data Portal. Reference labels for focal DNA amplifications were obtained from AmpliconRepository, which hosts outputs from the AmpliconSuite pipeline applied to whole genome sequencing (WGS) data [5, 12]. Amplification annotations included ecDNA, breakage–fusion–bridge, complex non-cyclic, and linear, as well as no-amplification states. These architectures are not mutually exclusive, and individual tumours may harbour multiple amplification patterns concurrently. The extent of their co-occurrence structure is illustrated in Figure 1b, with inset circles capturing additional pairwise overlaps that are visually compressed within the four-set diagram.

**Fig. 1:**
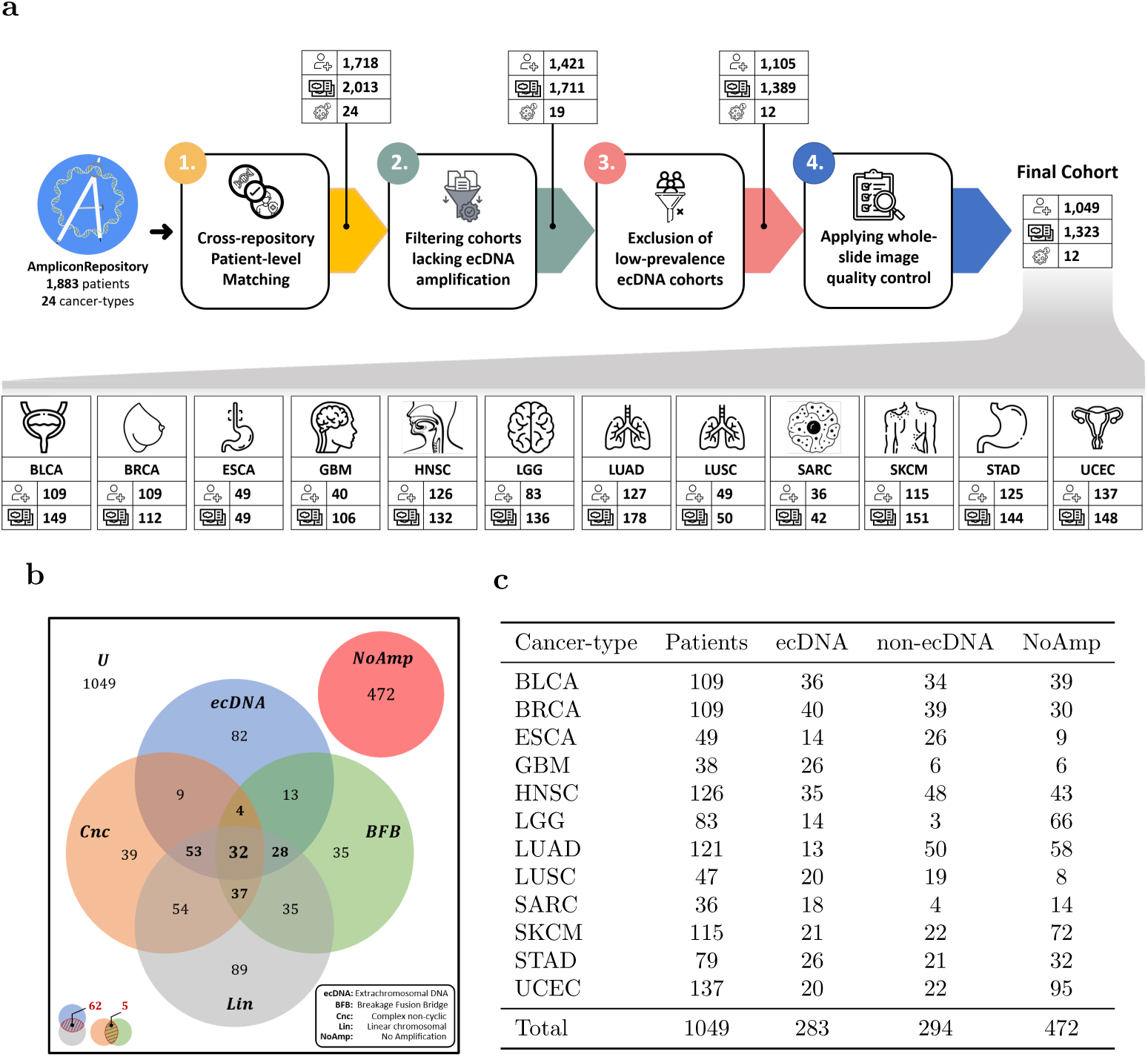
Cohort construction and genomic amplification landscape underlying the multi-cancer study. (a) Cohort assembly and data curation workflow. (b) Venn diagram illustrating co-occurrence of amplification architectures (extrachromosomal DNA(ecDNA), breakage–fusion–bridge events (BFB), complex non-cyclic amplifications (Cnc), and linear chromosomal amplifications (Lin)) among amplified tumours. (c) Final curated multi-cancer cohort after ecDNA-centric refinement and WSI quality control.

Because these sources differ substantially in cancer coverage, slide availability, and technical quality, cohort assembly required staged refinement. As outlined in Figure 1a, this process comprised cross-repository patient-level matching, tumour–level filtering based on amplification annotations, and slide-level quality control to remove corrupted or artefactual slides (details are provided in Methods). Following these steps, the final study population comprised 1,323 WSIs from 1,049 patients spanning 12 cancer types (Figure 1c). Each cancer type was analysed as a distinct sub-cohort to enable stratified evaluation while preserving a unified modelling framework. The resulting dataset exhibits considerable variation in tissue extent, histologic appearance, and class balance across tumour types, necessitating careful image preparation before model development.

## An end-to-end, slide-coherent MIL framework for routine H&E at diagnostic resolution

We developed AMIE (Augmented Multi-Instance learning with Interpretable attention), an end-to-end framework that infers slide-level ecDNA status directly from routine H&E WSIs under weak supervision. WSIs are first restricted to tissue-containing regions and tiled at 20× magnification into thousands of patches per slide; only slide-level labels are used, with no region annotations. AMIE jointly optimises a patch-level feature encoder and an attention-based pooling mechanism within a multiple-instance learning (MIL) framework that produces a compact slide representation and probability prediction. The attention weights provide post hoc localisation of morphology that contributes most strongly to the decision, supporting downstream biological interrogation.

To counter class imbalance, staining variability, and site-specific artifacts while preserving slide-level semantic coherence, we introduce coordinated slide-level augmentation strategies during training. These comprise three complementary families (Supplementary Figure A.1a): (i) Patch masking (constant, mean, or hybrid) stochastically obscures subsets of patches to prevent reliance on a few salient regions, promoting integration of distributed histological cues; (ii) Fourier-domain perturbations modulate low- and high-frequency components to diversify tissue architecture and texture without disrupting spatial layout; (iii) Stain-aware colour distortion (controlled adjustments to brightness, contrast, saturation, and hue with occasional greyscale conversion) reduce sensitivity to laboratory-specific appearance profiles. Ablation analyses demonstrate that these slide-coherent transformations mitigate overfitting and enhance cross-cancer generalisation (see Methods).

Unlike approaches that rely on frozen, precomputed embeddings, AMIE trains the encoder and MIL pooling mechanism jointly so that the ecDNA objective shapes patch-level representations. To support thousands of patches per slide at diagnostic resolution, we stream patch features and employ model-parallel training, preserving uninterrupted gradient propagation from slide-level predictions through the pooling operator into encoder parameters. This enables end-to-end learning at diagnostic resolution and improves discrimination relative to frozen-feature baselines. Moreover, the attention weights serve as a meaningful explanatory signal, facilitating interpretation of model focus in relation to underlying histomorphological patterns at diagnostic resolution. An overview of the AMIE framework is shown in Figure 2, and full implementation details, hyperparameters, and ablation protocols are provided in Methods.

**Fig. 2:**
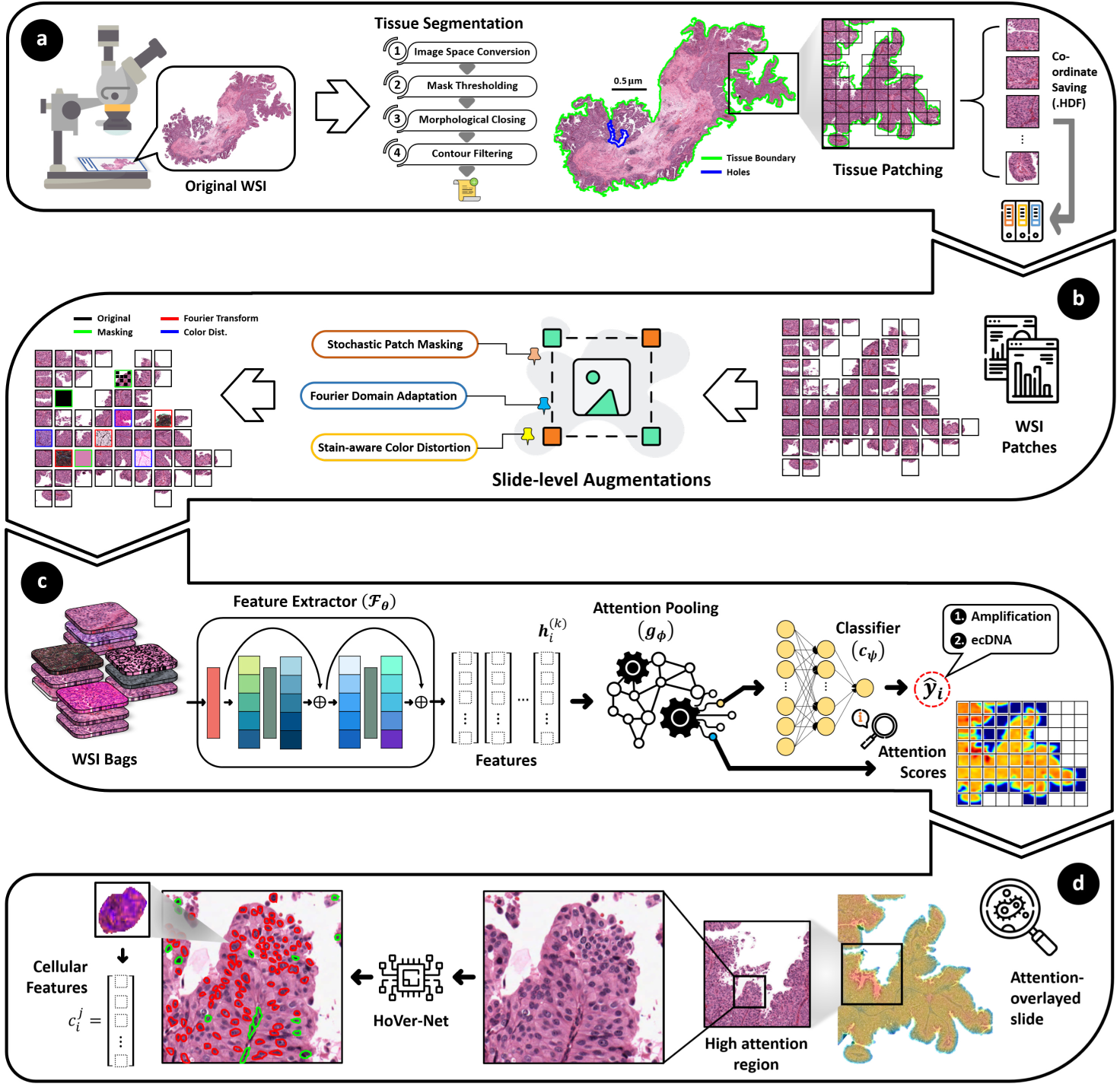
Overview of the AMIE framework for ecDNA-driven tumour detection from Whole-slide images. (a) WSIs are segmented to identify tissue regions and partitioned into fixed-size patches, forming slide-level bags under weak supervision. (b) Slide-level augmentation generates multiple coherent views of each slide by applying biologically motivated transformations across patch sets. (c) Augmented patch bags are encoded using a jointly trained ResNet-50 feature extractor and aggregated via attention-based pooling to produce slide-level predictions of amplification status. (d) Attention scores enable post hoc interpretability by localising highly informative regions, facilitating downstream cellular and morphological analysis.

## Genomic amplification leaves a detectable histologic signal

Before testing whether ecDNA-specific morphology is learnable, we first asked whether any genomic amplification leaves a detectable imprint in routine whole-slide histopathology images. Patients harbouring any focal amplification event (i.e., ecDNA, BFB, Cnc, or Lin) were labelled amplified, whereas those without such events were labelled non-amplified. As summarised in Figure 1b, this definition aggregates tumours across overlapping amplification architectures into a single positive class, thereby framing amplification detection as a broad, architecture-agnostic classification task. Labels were assigned at the patient level and inherited by all slides from that patient. To preserve disease-specific context, the framework was trained independently for each cancer type. Within each project, data were split into three patient-level, stratified folds with all slides from a given patient confined to a single split to prevent slide leakage across folds and evaluated using eight different performance metrics (see Methods) on held-out folds (Figure 3a).

**Fig. 3:**
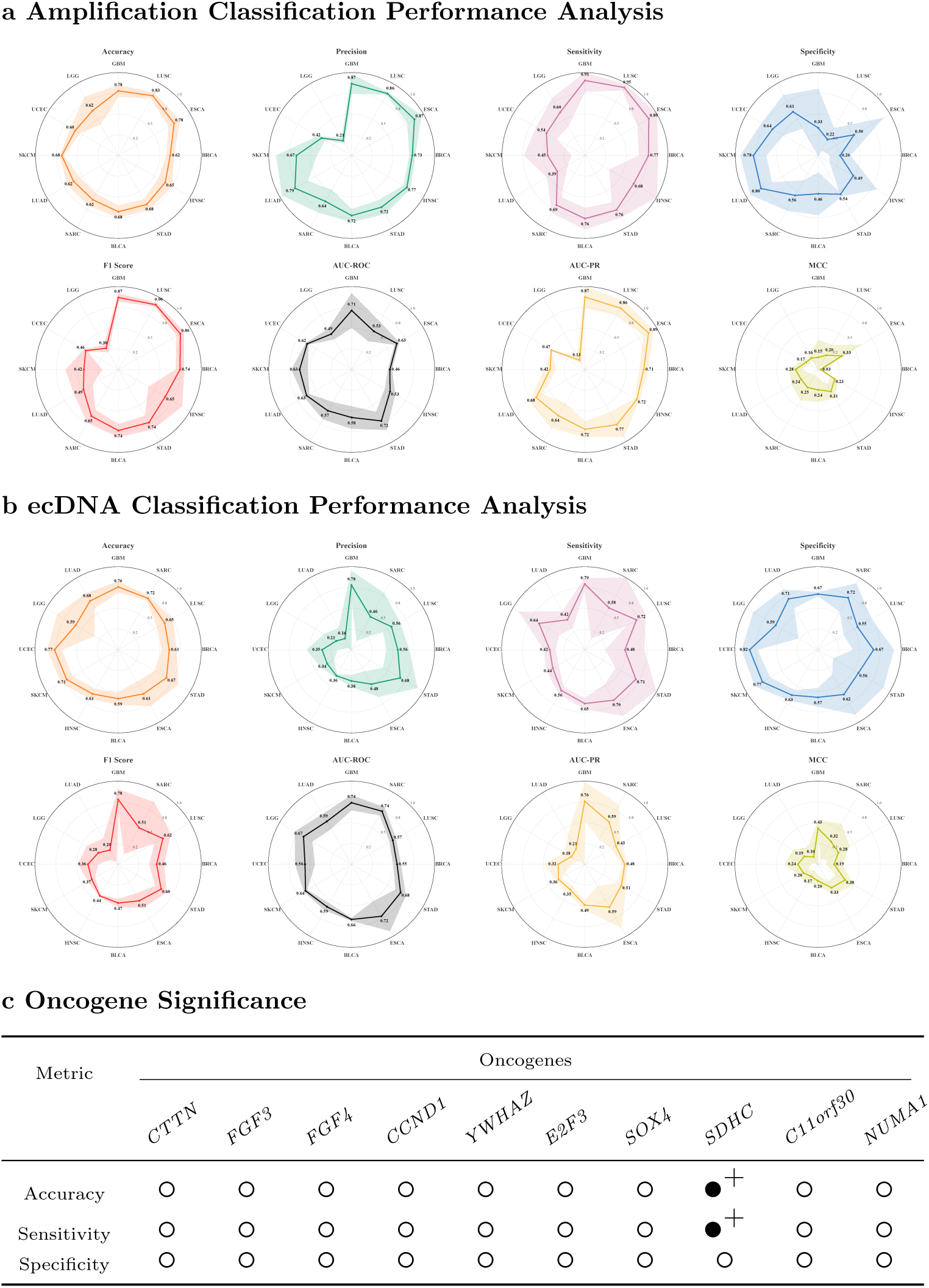
Cross-cancer performance of amplification and ecDNA classification with oncogene-stratified evaluation. (a) Genomic amplification detection. (b) ecDNA-specific classification. (c) Oncogene-stratified statistical comparison in BLCA. Filled symbols denote significant differences (*p <* 0.05), and polarity indicates direction of performance bias.

Cohorts with high amplification prevalence, including GBM, LUSC, and ESCA (proportion of tumours with amplifications: 0.82 − 0.85), showed high sensitivity and precision (GBM: sensitivity 0.91, precision 0.87; LUSC: 0.95, 0.86; ESCA: 0.89, 0.87) and strong AUC-PR (0.86 − 0.88), indicating reliable identification of amplified cases. Specificity, however, was low (0.22 − 0.50), and AUC-ROC was modest (0.53 − 0.71), reflecting the known behaviour of ROC metrics under imbalance and the model’s bias toward the majority positive class in these cohorts. BRCA (proportion 0.72) followed this pattern with a good F1-score (0.74) and AUC-PR (0.71) but low specificity (0.26) and low AUC-ROC (0.46).

Cohorts with intermediate amplification prevalence (0.52−0.65), e.g., HNSC, STAD, BLCA, and SARC, tended to show a more balanced sensitivity and specificity. STAD, in particular, combined moderate-to-high sensitivity (0.76) with higher specificity (0.54) and the best AUC-ROC in this set (0.72), alongside solid AUC-PR (0.77) and F1 (0.74), suggesting clearer separability of amplified vs. non-amplified morphology in gastric cancer. BLCA and SARC showed similar, balanced trends (F1-score ≈ 0.65−0.74; AUC-PR ≈ 0.64 − 0.72), whereas HNSC exhibited greater fold-to-fold variability (sensitivity 0.68 ± 0.34; specificity 0.49 ± 0.33), consistent with heterogeneous tissue patterns.

At lower prevalence, LUAD, SKCM, and UCEC (0.32 − 0.50) showed shifts in the precision-recall trade-off. LUAD (0.50) reached high precision (0.79) with reduced sensitivity (0.38) and higher specificity (0.80), indicating conservative positive calls. SKCM (0.33) presented moderate AUC-ROC (0.63) but a lower AUC-PR (0.42), closer to its prevalence baseline, and a reduced F1 (0.42). UCEC (0.32) demonstrated moderate discrimination (AUC-ROC 0.62) with balanced specificity (0.64) but lower precision (0.42) and modest AUC-PR (0.47), consistent with limited positive-class enrichment relative to its prevalence. LGG (0.13) approached chance levels on PR and ROC (AUC-PR 0.13, AUC-ROC 0.49) with low F1 (0.30), consistent with the scarcity of amplified cases and subtle morphological contrast.

Across 12 cancer cohorts, the framework achieved an overall accuracy of 0.66, precision of 0.62, sensitivity of 0.64, specificity of 0.64, F1 of 0.63, AUC-PR of 0.63, AUC-ROC of 0.68, and MCC of 0.28. In aggregate, these numbers indicate that amplification status leaves a learnable morphological signal in histopathology, with PR-family metrics (AUC-PR, F1) outperforming ROC under imbalance, and MCC confirming modest but genuine discrimination.

## ecDNA is distinguishable from other amplification contexts

To determine whether this approach can specifically distinguish ecDNA-amplified tumours from those with only chromosomal or without focal amplification, we redefined the classification task by labelling tumours with ecDNA amplification as ecDNA-positive and all others as ecDNA-negative. As illustrated in Figure 1b, ecDNA constitutes one component within a broader and overlapping amplification landscape, making this a more stringent discrimination problem than the amplification-versus–non-amplification setting. We applied the same per-cancer training protocol used for the amplification baseline: three-fold, patient-level stratified cross-validation, with patient-confined splits. Across cancer types, performance broadly tracked ecDNA prevalence and cohort size (see Figure 3b): cohorts with more ecDNA-positive patients tended to yield stronger PR-family metrics and MCC, whereas sparse-positive cohorts showed weaker and less stable signals. In higher-prevalence cancers (proportion ≥ 0.30; GBM, SARC, LUSC, BRCA, STAD), the model achieved a mean AUC-PR of 0.55 and an MCC of 0.32. GBM led this group (AUC-PR 0.76; MCC 0.43) with balanced precision and recall (0.78/0.79), aligning with genomic reports of frequent focal amplifications, chromosomal instability, and ecDNA-associated heterogeneity in glioblastoma [1, 18, 19]. STAD performed consistently (MCC 0.38), whereas LUSC favoured sensitivity over precision. Despite similar prevalence, SARC exhibited substantial fold-to-fold variability, likely reflecting limited training data (18 ecDNA-positive, 24 ecDNA-negative patients). BRCA was a notable exception: despite moderate prevalence (0.38) and a larger sample size (42 positive, 70 negative), discrimination was weaker (AUC-PR 0.48; MCC 0.19), suggesting that ecDNA-associated morphology in breast carcinoma may be subtler or more context-dependent. Intermediate-prevalence cohorts (ESCA, BLCA, HNSC; 0.27-0.29) showed moderate discrimination with increased variability, consistent with partial separability of ecDNA-positive foci within heterogeneous epithelium.

To further interrogate sources of misclassification, we examined per-cancer distributions of correct versus total predictions stratified by amplification context (Supplementary Figure A.2). For each tumour type, Venn diagrams partition patients into ecDNA-positive, non-ecDNA focal amplification, and no-amplification groups, reporting correct classifications within each category. In several cancers, including BRCA, LGG, and SARC, misclassification was not confined to sparse ecDNA cohorts but also occurred in tumours harbouring other focal amplifications. This pattern supports the hypothesis that overlapping genomic amplification architectures may introduce morphologic features that partially resemble ecDNA-driven patterns, thereby biasing predictions toward or away from the positive class. Conversely, in cancers such as GBM and STAD, correct classification rates were more consistently aligned with ecDNA status, suggesting clearer morphologic separation between amplification contexts.

At the lower end of the prevalence spectrum (SKCM, UCEC, LGG, LUAD; ≤ 0.18), class imbalance constrained positive-class learning. SKCM and UCEC reached high specificity (≥0.77) but modest MCC (≤0.24), indicating that apparent performance was partly driven by the dominant negative class. LUAD was most affected (prevalence 0.08; MCC 0.10), exhibiting a strong tendency toward majority-class predictions. LGG (prevalence 0.12) remained unstable across folds, consistent with few positive examples and subtle histological contrast.

Overall, across twelve cancers, the framework achieved an accuracy of 0.67 ± 0.02 and AUC-ROC 0.67 ± 0.00, with sensitivity 0.63 ± 0.04 and specificity of 0.68 ± 0.04. PR-focused metrics (AUC-PR 0.40 ± 0.12, F1 0.50 ± 0.01) and MCC (0.28 ± 0.01) indicate modest but consistent ecDNA signal capture beyond majority-class bias. Collectively, these findings suggest that ecDNA-associated morphology is recurrent yet cancer-type dependent. These trends emphasise prevalence-aware reporting and cohort-specific interpretation when deploying ecDNA predictors, providing a foundation for integrating molecular context into morphology-based prediction pipelines.

## Investigating the impact of amplified oncogenes on ecDNA predictions

Given that ecDNA often carries transcriptionally amplified oncogenes [6, 20], it remains unclear whether ecDNA predictions from histopathological images are influenced by specific oncogenic drivers or instead reflect oncogene-agnostic morphological features. Across cancer types, the most frequently amplified oncogenes exhibited heterogeneous enrichment between ecDNA- and chromosomal-amplified tumours. We therefore examined whether amplified oncogenes translate into systematic differences in prediction performance.

For each cancer type, we identified the top *k* = 10 most frequently amplified oncogenes. For each oncogene *O_k_*, patients were stratified into amplification-present (*O*^+^_*k*_) and amplification absent (*O*^−^_*k*_, potentially with other amplifications). Using out-of-fold predictions from the primary ecDNA classification framework, we compared subgroup accuracy, sensitivity, and specificity. Accuracy differences between *O*^+^_*k*_ and *O*^−^*_k_* were assessed using a two-sided Mann-Whitney U test on per-patient correctness indicators, whereas sensitivity and specificity differences were assessed using Fisher’s exact test on the corresponding contingency tables. To complement *p*-values, we also record the direction of any effect (better performance in *O*^+^_*k*_ vs. *O*^−^_*k*_) to flag potential subgroup biases even when absolute differences are modest (for details, see Methods).

In bladder cancer (BLCA), for example, some oncogenes were preferentially associated with chromosomal amplifications (e.g., *CTTN*), whereas others were enriched on ecDNA-amplifications (e.g., *YWHAZ*). Despite these genomic differences, model performance was largely comparable across amplification strata (Figure 3c). One exception was the tumour suppressor gene *SDHC*, a subunit of the succinate dehydrogenase enzyme, whose amplification coincided with higher accuracy and sensitivity without affecting specificity, indicating a context in which ecDNA predictions were modestly conditioned on specific amplified genes.

Extending this analysis across twelve cancer types revealed that oncogene-associated performance shifts were generally sparse and context-dependent. Several tumour types showed no detectable conditioning by oncogene amplification, whereas isolated effects were observed in others, including glioblastoma, lung adenocarcinoma, and lung squamous cell carcinoma. A pooled pan-cancer analysis further indicated that only a small subset of oncogenes exhibited consistent associations with prediction behaviour across tissues, highlighting the potential for cancer-type composition to confound gene-specific effects. Overall, these findings suggest that ecDNA prediction from routine histopathology is broadly robust to the identity of amplified oncogenes. Rather than being driven by individual oncogenic events, model performance appears to reflect more general morphological correlates of ecDNA, supporting the biological specificity of the approach while motivating stratified reporting to capture rare, context-specific deviations. Full per-cancer and pooled analyses are provided in Supplementary Figures A.4–A.16.

## End-to-end learning outperforms frozen foundation embeddings

We hypothesized that genomic amplification driven by ecDNA manifests as a biologically subtle and spatially heterogeneous signal in histopathology. It remains unclear whether such signals can be adequately captured using generic or frozen feature representations, including those derived from large-scale histopathology foundation models. We therefore asked whether effective discrimination would instead require feature spaces that are explicitly shaped by slide-level supervision and the downstream classification objective. To address this, we compared our end-to-end trained feature extractor against widely used pretrained representations, including ImageNet-based features and three histopathology-specific foundation models (Virchow [21], UNI [22], and CTransPath [23]). In all cases, pretrained embeddings are kept frozen, and only the downstream components of the framework are trained. All models are evaluated under identical training and evaluation protocols, with performance averaged across 12 cancer types using three-fold cross-validation (Figure 4a).

**Fig. 4:**
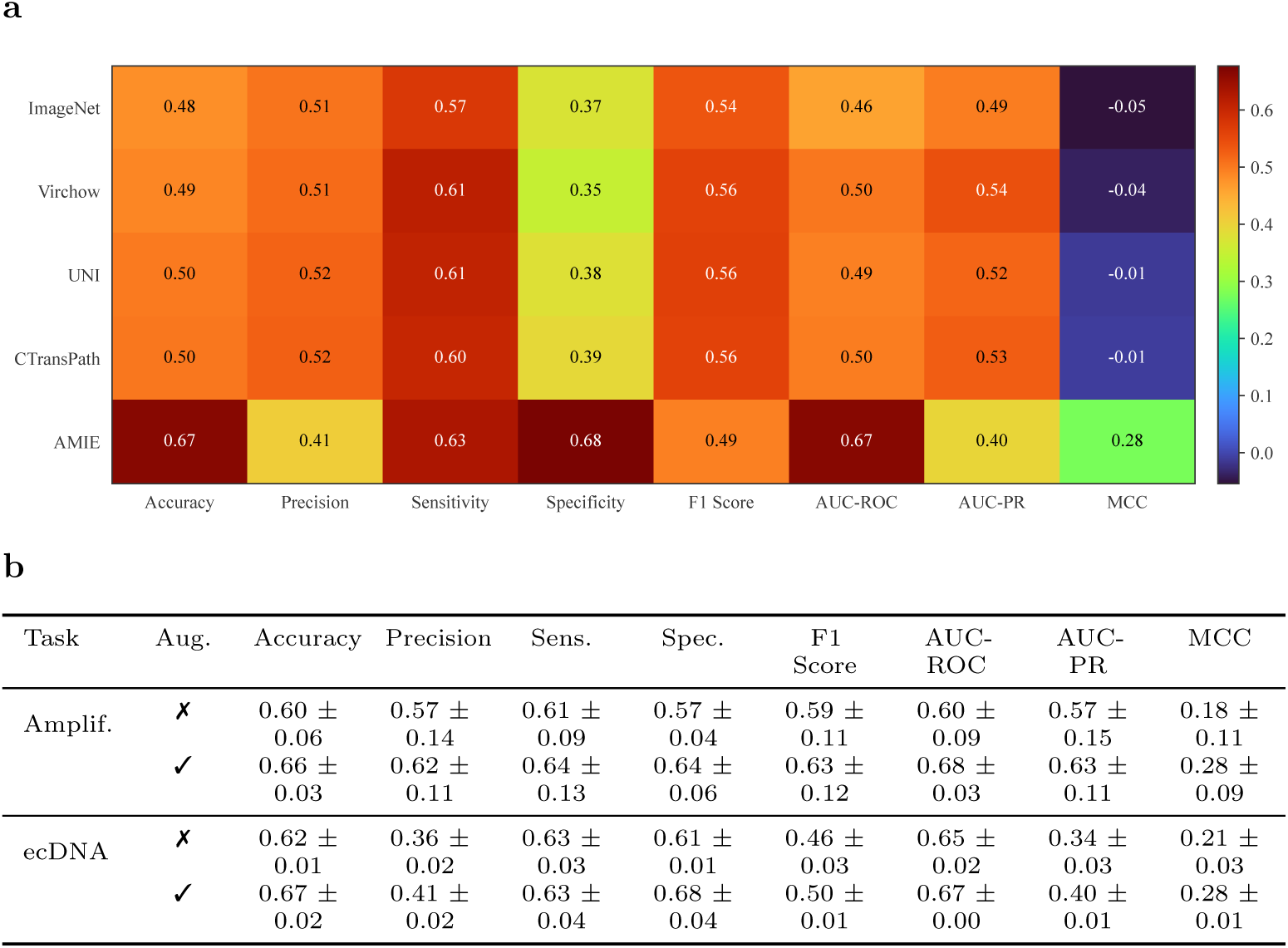
Impact of feature representation and slide-level augmentation on histopathology-based amplification classification. (a) Cross-cancer performance comparison between end-to-end training and frozen foundation-model features for ecDNA classification. (b) Mean cross-cancer performance with and without coordinated slide-level augmentation for amplification and ecDNA tasks.

Across all evaluated pretrained representations, discrimination of ecDNA-driven amplification status remained limited. ImageNet-based features exhibited near-random behaviour, with AUC-ROC values below 0.45 and a negative MCC, indicating systematic misalignment between the learned feature space and the underlying biological signal, despite moderate sensitivity. Histopathology-specific foundation models provided modest improvements in probabilistic discrimination, but these gains did not translate into robust, balanced classification. Across Virchow, CTransPath, and UNI embeddings, AUC–ROC values clustered narrowly around 0.48 − 0.51, while MCC remained close to zero, reflecting an inability to jointly control false positives and false negatives under pronounced class imbalance. Even UNI, which performed best among frozen representations, achieved only marginally positive MCC, indicating weak separation between amplified and non-amplified tumours.

In contrast, end-to-end feature learning led to a marked and consistent improvement across clinically relevant metrics. Relative to the strongest frozen representation, the proposed approach increased AUC-ROC by approximately 0.17 and improved MCC by more than an order of magnitude. Notably, this gain is driven primarily by a pronounced increase in specificity (0.68 versus ≤0.39 for all pretrained models), indicating improved suppression of false-positive predictions in non-amplified tumours, while sensitivity remained comparable. This shift suggests that end-to-end training enables the model to learn amplification-relevant visual cues that are systematically missed by task-agnostic representations, rather than merely re-weighting decision thresholds. Together, these results demonstrate that coupling representation learning with slide-level supervision is critical for shaping feature spaces that capture this biological signal, leading to substantial gains in discrimination and balanced classification performance.

## Slide-level augmentation improves generalisation and calibration

To systematically assess the impact of slide-level augmentation, we first conducted a controlled evaluation on the TCGA-BLCA cohort using a fixed train-test split (96 training and 45 test slides; 27/12 ecDNA-positive and 69/33 ecDNA-negative cases, respectively). Under identical architectural and optimisation settings, separate models are trained using a single augmentation strategy at a time and compared against a non-augmented baseline. Performance across configurations is summarised in Supplementary Table A.1, from which the highest-ranking setting in each augmentation family was selected for further analysis.

Augmentation consistently improved performance relative to baseline, with the largest gains observed in metrics capturing balanced error handling and minority-class recognition. In particular, stochastic patch masking and stain-aware colour produced the strongest improvements by increasing MCC (mean absolute gain: +0.48) and AUC-PR (mean gain: +0.21), indicating enhanced discrimination that is not driven solely by specificity. Although both strategies achieved comparable overall accuracy, they differed in their operating characteristics: patch masking favoured higher specificity (0.97) with moderate sensitivity (0.42), whereas colour distortion achieved higher sensitivity (0.50) at a modest reduction in specificity (0.94). On the other hand, Fourier domain adaptation also improved specificity (0.97) and AUC-PR (0.60) relative to baseline but exhibited limited sensitivity (0.25), suggesting that frequency-based perturbations primarily enhance probability ranking and false-positive control rather than minority-class recall. Based on these characteristics, patch masking and colour distortion emerged as the most effective augmentation strategies, while retaining Fourier domain adaptation as a complementary option.

Guided by these BLCA-based findings, we fixed the highest-ranking configurations from each family and extended the evaluation to all 12 cancer types under identical training and evaluation protocols. Figure 4b summarises the aggregated performance across all tumour types, with metrics reported as mean±standard deviation over three-fold cross-validation. Across cancer types, augmentation conferred consistent benefits, although the pattern differed between tasks. For amplification-driven tumour prediction, augmentation improved error balance and recall, yielding a moderate but consistent increase in MCC (≈ +0.10) and sensitivity (≈ +0.03), while precision and AUC-ROC showed modest improvements (≈ +0.05 and ≈ +0.08, respectively). In contrast, ecDNA-driven tumour classification exhibited broader gains in discrimination and calibration, with consistent improvements in accuracy (≈ +0.05), specificity (≈ +0.07), AUC-ROC (≈ +0.02), and AUC-PR (≈ +0.06). These coordinated gains translated into an MCC improvement of approximately +0.07 (≈ 33% relative to baseline), without a corresponding loss in sensitivity.

Taken together, these results demonstrate that slide-level augmentation improves performance across heterogeneous cancer cohorts, but with task-specific effects. Amplification prediction primarily benefits from improved recall and error balance, whereas ecDNA classification shows stronger gains in probabilistic discrimination and specificity. Detailed analyses of individual augmentation configurations, the ranking procedure, and the full cancer-specific results are provided in Methods.

## Interpretable attention aligns with ecDNA-associated nuclear anomalies in GBM

To explore the morphological evidence underlying ecDNA prediction in AMIE, we conducted a nucleus-level anomaly-detection study in the GBM cohort. We focus on nuclei because ecDNA is a chromatinised DNA element residing in the nucleus, and any H&E-detectable morphological footprint is therefore most likely to manifest at the nuclear level.

All nuclei were segmented using HoVer-Net [24], which provides instance-level nuclear masks and coarse cell-type labels but does not perform anomaly detection or feature learning. From each segmented nucleus, we subsequently extracted a rich set of handcrafted features capturing nuclear morphology, colour statistics, hematoxylin and eosin optical density, and chromatin texture. In total, this process yielded over 29 million nuclei across the cohort. Based on slide-level ecDNA annotations, nuclei were grouped into three populations: Pop1, GBM without ecDNA or focal amplifications (14 WSIs; 8.98 M nuclei); Pop2, GBM with focal amplifications but no ecDNA (33 WSIs; 4.89 M nuclei); and Pop3, ecDNA-positive GBM (64 WSIs; 16.01 M nuclei). Each nucleus was associated with the AMIE attention score of the patch in which it was embedded, enabling spatial alignment between model attention and nucleus-level morphology.

We first qualitatively assessed the spatial relationship between AMIE attention and nuclear anomalies. In ecDNA-positive slides, nuclei identified as anomalous by an Isolation Forest tended to co-localise with regions receiving high AMIE attention (Figure 5b). Importantly, this correspondence is subtle and not driven by a visually obvious increase in the absolute number of anomalous nuclei. Instead, anomalous nuclei remain sparse and are distributed across both high- and low-attention patches. Representative highest- and lowest-attention patches further illustrate this behaviour (Figure 5c). The top example shows an ecDNA-positive GBM slide in which high-attention patches contain sparse morphologically anomalous neoplastic nuclei (cyan) interspersed among otherwise typical tumour cells. While the absolute frequency of anomalous nuclei appears comparable between high- and low-attention patches, high-attention regions exhibit subtle but consistent shifts in nuclear appearance and chromatin organisation. In contrast, the ecDNA-negative (focal-amplification) example shows differences in tissue composition between high- and low-attention patches but lacks comparable anomalous nuclear patterns. In both cases, patches are predominantly composed of neoplastic nuclei, indicating that AMIE attention is not driven solely by tumour cell density.

**Fig. 5:**
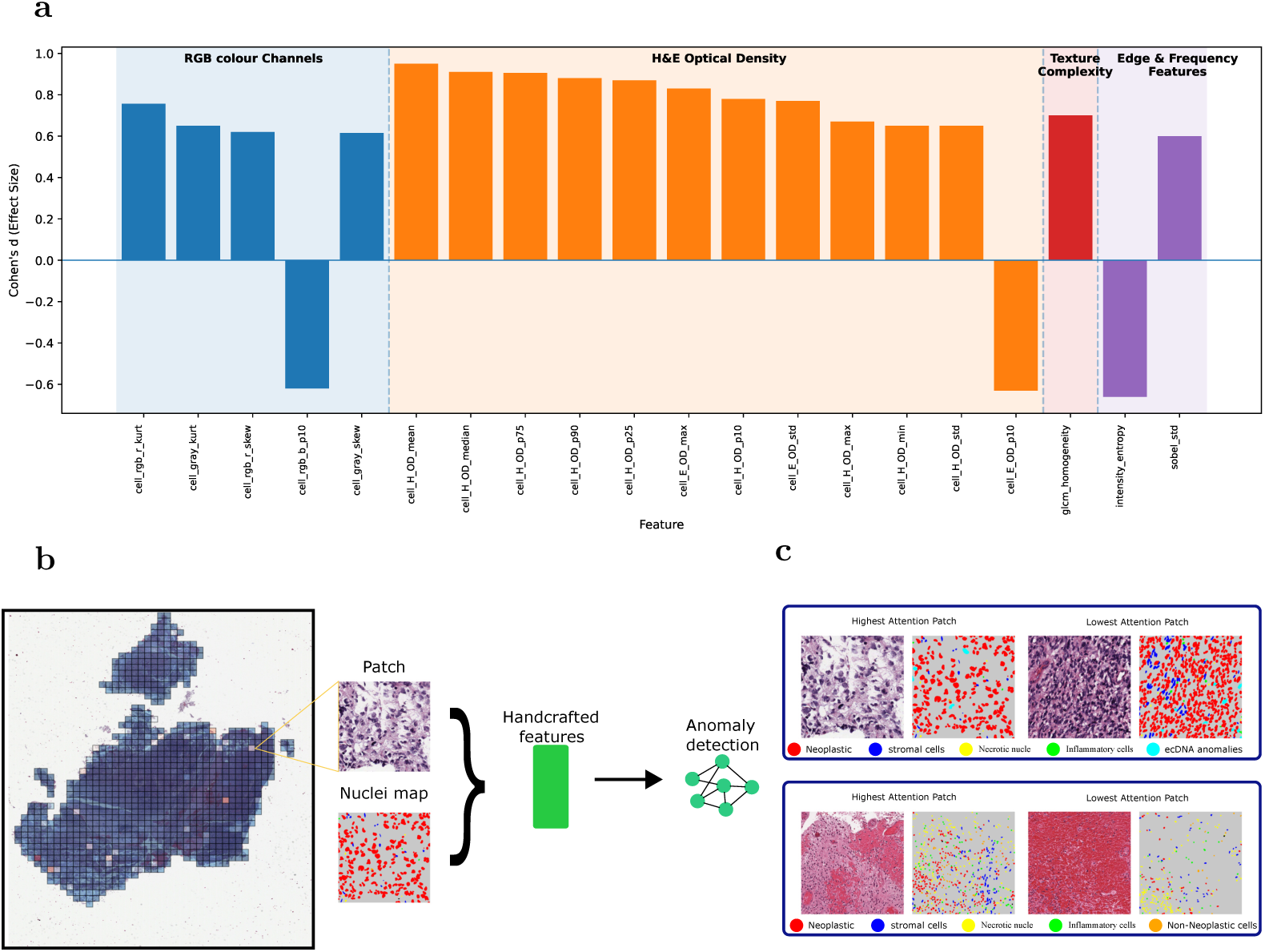
Multi-scale explainability of AMIE ecDNA predictions in GBM. (a) Top discriminative nuclear features separating anomalous from non-anomalous Pop3 nuclei, quantified by Cohen’s *d*. (b) Qualitative alignment between AMIE attention and nucleus-level anomaly detection in a representative ecDNA-positive WSI, showing patch tiling, nuclei segmentation, and anomalous nuclei identified by Isolation Forest. (c) Representative patches with the highest and lowest AMIE attention scores.

To quantify these observations, we framed the problem as rare-event detection, reflecting uncertainty as to whether ecDNA-associated morphology is widespread or confined to rare cellular subpopulations. An Isolation Forest was trained on nuclei from Pop1 and Pop2 (13.9 M nuclei) to learn a baseline model of canonical GBM nuclear morphology, using the full feature set and a conservative contamination rate of 0.1%.

When applied to Pop3, the model identified 160,146 nuclei (1.0%) as morphologically anomalous. Feature-level comparison of anomalous versus non-anomalous Pop3 nuclei using Cohen’s *d* (Figure 5a) showed that the strongest shifts were driven by hematoxylin optical-density statistics, followed by chromatin texture descriptors such as GLCM homogeneity, with smaller contributions from edge- and colour-related features. Positive effect sizes indicate increased nuclear staining intensity and more homogeneous chromatin texture among anomalous nuclei, consistent with altered chromatin organisation.

Although this analysis reveals an interpretable morphological signal in ecDNA-positive GBM, the Isolation Forest is limited as a standalone approach: the detected fraction is constrained by a predefined contamination rate, and the model operates at the single-nucleus level without spatial context. In contrast, AMIE integrates signals from thousands of nuclei at the patch and slide levels and can capture weaker, distributed patterns across spatial scales. Together, these results indicate that ecDNA-associated morphology in GBM is rare, heterogeneous, and spatially structured, necessitating multi-instance, context-aware models such as AMIE for reliable detection.

## AMIE predicted ecDNA status recapitulates adverse prognosis

As ecDNA has previously been shown to associate with poor survival in the TCGA cohort [5], we examined whether imaging-based predicted ecDNA status similarly captures survival-related biology. As a reference, we first evaluated ecDNA status determined by WGS within our current cohort. Consistent with previous findings, AmpliconArchitect-classified ecDNA was associated with significantly shorter five-year overall survival (*P=0.012* by log-rank test) (Figure 6a). However, in contrast, our analysis was limited to a smaller patient subset and compared ecDNA-positive tumours against both tumours harbouring no amplifications and those harbouring chromosomal amplifications. Within this cohort, the survival association did not remain significant after stratifying by tumour type (*HR=1.12, 95%CI 0.9-1.4, P=0.31*). Importantly, a significant survival difference between patients whose tumours harbour ecDNA against patients whose tumours contain chromosomal but no ecDNA amplifications has only recently been demonstrated in larger adult and paediatric cancer patient populations [6, 8].

**Fig. 6:**
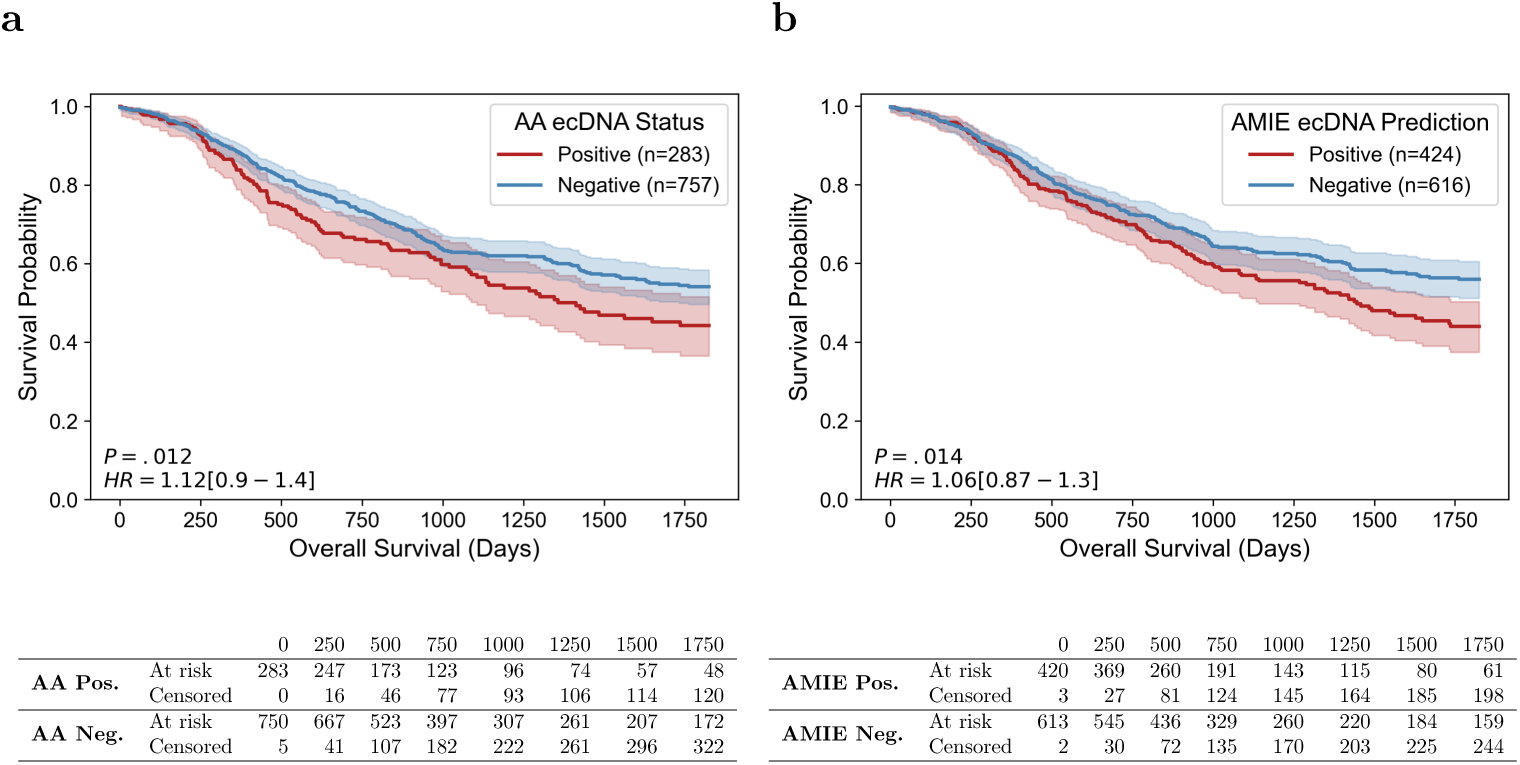
Association between ecDNA status and overall survival. Kaplan-Meier curves comparing five-year overall survival, stratified by ecDNA status derived from (a) AmpliconArchitect (AA) or (b) AMIE. P-values are from log-rank tests. Hazard ratios with 95% CIs were estimated from Cox proportional hazard models controlling for tumour type, with ecDNA-negative as reference. Shaded regions represent 95% CIs.

AMIE-predicted ecDNA likewise showed a significant association with overall survival (*P=0.014* by log-rank test), closely mirroring the effect observed with WGS within the same cohort (Figure 6b). As with the AmpliconArchitect-based comparison, this association did not persist when controlling for tumour type *HR=1.06, 95%CI 0.87-1.3, P=0.55*, a pattern consistent with the reduced sample size and the more conservative comparison group that includes tumours with chromosomal but without ecDNA amplifications used in the present study. Together, these results demonstrate that our purely imaging-based method recapitulates the survival patterns observed with DNA sequencing-based methods when evaluated under identical cohort constraints.

## Discussion

This study shows that routine H&E histopathology contains a recoverable signal of ecDNA, enabling the prediction of ecDNA status across multiple tumour types. The central contribution is to demonstrate that a morphology-only model can capture correlates of ecDNA, an inherently molecular hallmark, at the slide level. That performance remains broadly stable across amplified oncogene contexts suggests the model leverages general morphologic consequences of ecDNA (potentially proliferation dynamics, nuclear atypia, necrosis patterns, stromal organisation) rather than proxying a small set of driver events.

Robust evaluation is challenging because ecDNA prevalence varies widely by cancer type, creating substantial class imbalance. Threshold-independent and threshold-dependent metrics were therefore considered jointly (including AUC-PR, F1, and MCC alongside AUC-ROC, accuracy, sensitivity, and specificity) to avoid overstating performance through an abundance of true negatives in ecDNA-rare settings [25]. Oncogene-conditioned analyses further indicated that performance shifts are sparse and context-specific, reinforcing the value of stratified reporting to surface pockets of degradation or unexpected gains that may matter clinically.

Label quality is a primary limitation. Ground truth depends on short-read reconstruction pipelines in which AmpliconArchitect (AA) is commonly used; AA’s non-trivial error rate of 10-15%, spanning false positives (e.g., mislabelling BFB-like or complex linear amplicons as circular) and false negatives (missed ecDNA when breakpoints are under-resolved), introduces supervision noise. Such noise likely depresses apparent headroom, blurs decision boundaries, and can manifest as site- or subtype-specific errors attributed to the model. Consistent cross-validation performance and limited gene-conditioned drift suggest that the histologic signal exceeds this noise floor, yet definitive assessment will require orthogonal validation (e.g., long-read sequencing, optical mapping, or ecDNA-targeted FISH) and, where possible, re-training with adjudicated labels to quantify gains achievable with improved supervision.

Generalisability and clinical readiness remain to be established. TCGA slides offer diversity in staining and scanning but are retrospective and research-grade. Domain shifts, pre-analytic variation, scanner differences, necrosis burden, and site-specific protocols can degrade performance. Prospective, multi-centre studies with locked models, pre-registered analyses, and calibration assessments are needed to determine real-world operating points and the clinical consequences of false positives and false negatives. In near-term practice, an H&E-based predictor is best positioned as a triage tool: prioritizing samples for confirmatory sequencing, enriching trials for ecDNA-positive disease, and flagging cases for intensified monitoring where ecDNA portends aggressive behaviour and resistance.

In sum, ecDNA appears to imprint a histomorphological phenotype that is recoverable from standard slides. Slide-based inference offers a scalable, low-cost complement to sequencing, contingent on rigorous external validation, improved ecDNA ground truth, and transparent, stratified reporting to manage risk at deployment.

## Supporting information

amplification-driven and ecDNA-driven prediction tasks in the Supplementary Material.

amplification-driven and ecDNA-driven prediction tasks in the Supplementary Material.

amplification-driven and ecDNA-driven prediction tasks in the Supplementary Material.

## Acknowledgment

This work is supported by a generous endowment from the Clayes Foundation to the Research Center for Neuro-Oncology and Genomics within the Rady Children’s Institute for Genomic Medicine, a Hannah’s Heroes St. Baldrick’s Scholar Award (L.C.), the Dragon Master Foundation (L.C.), funding from the National Institutes of Health (NIH) National Institute of Neurological Disorders and Stroke Institute R01NS132780 (L.C.), and the NIH National Cancer Institute P30CA030199 (L.C.), P30CA030199-42-S1 (L.C.). Support through grant P30CA030199 to the Genomics core facility at Sanford Burnham Prebys (NCI designated Cancer Center) is gratefully acknowledged. The content is solely the responsibility of the authors and does not necessarily represent the official views of the National Institutes of Health.

The authors also acknowledge the support of the Ministry of Science and Culture of Lower Saxony through funds from the program zukunft.niedersachsen of the Volkswagen Foundation for the ‘CAIMed – Lower Saxony Center for Artificial Intelligence and Causal Methods in Medicine’ project (grant no. ZN4257). The authors acknowledge Hannover Medical School for providing MHH-HPC resources and technical support that have contributed to the research results reported within this paper.

## Methods

### Extended Description of Cohort Assembly and Refinement

This section provides a detailed description of the stepwise cohort assembly and refinement process. Starting from the integration of TCGA whole-slide histopathology images with focal amplification annotations from the AmpliconRepository, we outline the successive patient-level matching, ecDNA-centric cohort filtering, and slide quality-control steps that resulted in the final curated multi-cancer cohort used in this study.

#### Cross-repository patient-level matching

Histopathology and genomic data were available at different scales across the two sources. TCGA contributed 11,765 diagnostic H&E-stained whole-slide images from 9,640 patients spanning 33 cancer types, whereas the AmpliconRepository provided focal DNA amplification annotations for 1,883 patients across 24 cancer types. To align these datasets, patients from the AmpliconRepository were matched to TCGA cases using unique patient identifiers. Only patients for whom both histopathologic WSIs and focal amplification annotations were available were retained for downstream analysis. Amplification labels included ecDNA, breakage–fusion–bridge, complex non-cyclic, linear, and no-amplification states, as defined by the AmpliconSuite pipeline. Following patient-level matching, this step yielded 1,718 patients with 2,013 WSIs across 24 cancer types.

#### Filtering cohorts lacking ecDNA amplification calls

Next, we excluded cancer types for which no ecDNA-positive cases were present in the matched cohort. Although both amplification-driven and ecDNA-driven classification settings were evaluated in this study, identifying histomorphologic correlates of ecDNA-driven tumour amplification directly from WSIs constituted the primary focus. Cancer types lacking ecDNA events cannot support this objective and were therefore excluded at this stage. Based on this criterion, five cancer types (COAD, KIRC, READ, THCA, and UVM) were removed from further analysis. After this filtering step, the cohort comprised 1,421 patients with 1,711 WSIs across 19 cancer types.

#### Excluding low ecDNA-prevalence cohorts

To ensure reliable training and evaluation within each cancer type, we further excluded cohorts with very low numbers of ecDNA-positive cases. Under the three-fold cross-validation protocol used throughout this study, cancer types with fewer than 10 ecDNA-positive patients would yield at most three positive samples in each test fold. At this scale, performance estimates become highly sensitive to single-sample variation, leading to unstable and potentially misleading assessments of model behaviour. Based on this criterion, seven cancer types (CESC, DLBC, KICH, KIRP, LIHC, OV, and PRAD) were excluded due to insufficient ecDNA-positive sample counts. After this refinement step, the cohort comprised 1,105 patients with 1,389 WSIs across 12 cancer types.

#### Whole-slide image quality control

All remaining WSIs underwent a hybrid (manual + automated) quality control to identify technical artifacts that could confound computational histopathologic analysis. Slides were excluded if they were corrupted, lacked discernible tissue regions, or contained extensive pen markings or other non-biological artifacts. This quality-control step removed 66 WSIs from 56 patients, representing a substantial source of potential noise that is often under-reported in large-scale computational pathology studies, including analyses of TCGA lung cancer cohorts. Representative examples of excluded slides are shown in Supplementary Figure A.1b.

The final cohort composition is summarised in Figure 1c. Patients are stratified into three biologically meaningful groups: those exhibiting ecDNA amplification (including cases co-occurring with other amplification architectures), those exhibiting non-ecDNA focal amplifications (e.g., BFB, linear, or cyclic architectures without ecDNA), and those lacking focal amplification events.

This stratification directly supports both downstream classification settings used in this study. For amplification-driven classification, ecDNA and non-ecDNA cases are treated as amplified positives against non-amplified negatives. For ecDNA-driven classification, ecDNA-positive cases constitute the positive class, while non-ecDNA and non-amplified cases together form the negative class.

### Whole-slide Image Processing

WSIs are high-resolution tissue scans that capture diagnostically relevant tumour regions but also contain substantial non-informative content, such as background whitespace, pen markings, and preparation artifacts. To restrict model inputs to informative tissue while maintaining computational efficiency, we implemented a modular slide-processing pipeline that identifies tissue regions and subdivides them into fixed-size image patches. Our approach builds on the CLAM framework [26] with targeted refinements to improve robustness across diverse tissue morphologies, cancer types, and slide preparations.

#### Tissue segmentation

Tissue-containing regions are identified using a classical image-processing approach designed to separate stained tissue from background and slide artifacts. This step is particularly important for TCGA slides, where tissue coverage, cellularity, and scanner-specific artifacts vary substantially across cancer types and institutions. As illustrated in Figure 2 (a), each WSI is first read at a downsampled resolution (typically 32×) to reduce processing burden while preserving global tissue structure. The image is converted from RGB to hue–saturation–value (HSV) color space, and the saturation channel is smoothed using a median filter to emphasise stained tissue regions. A binary tissue mask is then generated by thresholding the smoothed saturation channel, using either Otsu’s method [27] or empirically fixed thresholds depending on slide-specific staining variability. To further refine the mask, morphological closing [28] is applied to connect fragmented tissue regions and remove small internal holes. Finally, candidate tissue regions are identified via contour detection [29] and filtered based on an area threshold to exclude spurious components. The resulting tissue mask delineates contiguous tissue regions and is used to constrain subsequent patch extraction to tissue-containing areas only.

#### Patch generation

Following tissue segmentation, WSIs are subdivided into smaller image patches to support scalable processing and learning. A regular coordinate grid is constructed across each WSI at a predefined magnification level, and candidate patch locations are filtered using the tissue segmentation masks to retain only coordinates within tissue-containing regions and outside any enclosed holes. For each retained coordinate, a fixed-size image patch is extracted from the original WSI. Patch extraction parameters, including magnification level, size, overlap, and spatial coordinates, are recorded and stored in a hierarchical data format (HDF5) to facilitate downstream reproducibility.

### Detailed Evaluation of Slide-level Augmentation Strategies

While the main manuscript reports only the performance comparison with and without employing augmentations, the analyses here systematically examine the behaviour of multiple parametrisations within each family and describe the metric-agnostic ranking framework used to identify robust augmentation settings. Together, these results provide quantitative justification for the augmentation choices adopted in the primary experiments.

Augmentation strategies are evaluated on the TCGA-BLCA cohort using identical data splits, model architectures, and optimisation settings to ensure direct comparability across configurations. Model performance is assessed using the same evaluation metrics reported throughout the study. Table A.1 summarises the held-out test-set performance of all evaluated augmentation configurations across the three augmentation families, alongside the non-augmented baseline.

#### Family-wise performance trends across augmentation configurations

Across all augmentation families, substantial heterogeneity can be observed in both absolute performance and metric-specific behaviour. Compared to the non-augmented baseline, most augmentation strategies improved balanced performance metrics such as MCC and F1 score; however, the magnitude, stability, and metric trade-offs of these gains varied markedly across parametrizations.

Within the stochastic patch masking family, simple masking strategies such as constant-value or mean-value replacement yielded moderate improvements over the baseline, particularly in sensitivity and AUC-based metrics. These gains, however, are accompanied by only modest increases in MCC (≤ 0.29), indicating limited overall class separation. In contrast, the hybrid masking strategy consistently outperformed other masking configurations across most metrics, achieving an MCC of 0.50 compared to 0.03 without augmentation, while maintaining high specificity (0.97). This pattern suggests that combining multiple masking priors provides stronger regularisation without suppressing discriminative tissue features. Conversely, aggressive blur-based masking markedly reduced sensitivity (0.25), indicating that excessive attenuation of local morphology can impair detection of ecDNA-associated patterns.

A distinct but complementary pattern emerged for Fourier domain adaptation, where performance is strongly dependent on frequency selection. While low-pass filtering improved sensitivity relative to the baseline, it yielded modest class separation overall. High-pass filtering, by contrast, substantially increased specificity (0.97) and precision (0.75), but at the cost of reduced sensitivity (0.25), reflecting a tendency toward confident majority-class predictions. These results suggest that frequency-selective perturbations can amplify certain textural cues while simultaneously suppressing information relevant for detecting ecDNA-positive cases.

Stain-aware colour distortions demonstrated the greatest variability across configurations. Deterministic transformations, such as isolated Gaussian blurring in stain space, collapsed predictive performance entirely (*F* 1 = 0), underscoring the risk of over-regularisation when stain perturbations are applied without sufficient variability. In contrast, randomised colour distortions, particularly when combined with greyscale conversion, yielded consistent improvements across multiple metrics, achieving MCC values exceeding 0.50 and balanced gains in both sensitivity and specificity. These findings indicate that stochastic colour variation better approximates real-world staining heterogeneity without erasing diagnostically relevant morphology.

Taken together, these family-wise analyses reveal a consistent trend: augmentation configurations that introduce controlled stochastic and parameter diversity tend to outperform both deterministic transformations and the non-augmented baseline. In contrast, overly aggressive or narrowly parametrised augmentations frequently degrade performance, emphasising the importance of balancing regularisation strength with preservation of discriminative histologic signals.

#### Metric-agnostic ranking-based selection of augmentation parameters

Because different augmentation strategies exhibited divergent behaviour across evaluation metrics, we avoided selecting configurations based on any single metric. Instead, we adopted a ranking-based aggregation scheme to identify parametrisations that demonstrated consistently strong performance across complementary criteria. For each augmentation family, configurations are independently ranked for each of the eight evaluation metrics, with rank 1 assigned to the best-performing configuration and rank N to the weakest, where N denotes the number of configurations evaluated within that family. These metric-specific ranks are then averaged to obtain a single summary score per configuration, reflecting overall robustness rather than peak performance on any individual metric. Configurations with the lowest average rank are selected as the representative settings reported in the main text.

#### Cancer–specific performance with and without augmentation

To assess the consistency of augmentation effects beyond the TCGA-BLCA cohort, we reported full cancer-type–specific performance comparing augmented and non-augmented models, amplification-driven and ecDNA-driven prediction tasks in the Supplementary Material. While the magnitude of improvement varies across tumour types, augmentation consistently improves balanced metrics such as MCC and AUC-PR in the majority of cohorts. These findings support the use of slide-level augmentation as a generalisable strategy rather than a BLCA-specific optimisation.

### End-to-end multiple instance learning architecture

Most histopathology-based MIL frameworks adopt a decoupled training strategy in which patch-level features are pre-trained and kept fixed, while only the slide-level aggregation module is optimised [26, 30]. We instead implement a fully end-to-end MIL framework in which the patch encoder, pooling operator, and classifier are jointly optimised under slide-level supervision. For a given slide *s_i_*, we extract a set of *N_i_* patches {*x*^(^*^i^*^)^}*^Ni^* . Each patch *x*^(^*^i^*^)^ is encoded by a convolutional network *f_θ_*, yielding an embedding *h*^(^*^i^*^)^ ∈ R*^d^*. All encoder parameters are updated jointly with the slide-level objective, allowing instance representations to be shaped directly by the amplification classification task rather than reflecting generic histological features learned from external data. To aggregate patch embeddings into a slide-level representation, we employ gated attention pooling, a learnable permutation-invariant operator that assigns an importance weight to each patch. For slide *s_i_*, attention weights are computed as

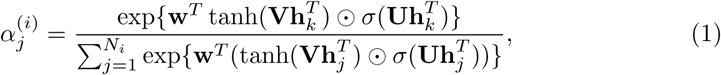

where V and U are learnable projection matrices, and ⊙ denotes element-wise multiplication. The slide representation is then obtained as

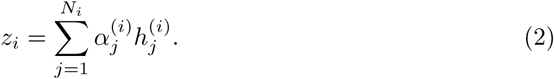

Because attention weights are computed at the patch level, the framework can optionally return attention scores alongside slide-level predictions, enabling post hoc visualisation of high-attention regions; however, attention weights should not be interpreted as causal attribution. The pooled representation *z_i_* is passed through a linear classifier to produce the slide-level logit *y*^*_i_*. In the binary setting, logits are converted to probabilities via a sigmoid function during evaluation, while training operates directly on logits. To account for class imbalance, we optimise a weighted binary cross-entropy loss. Given predicted probability *y*^*_i_* ∈ [0, 1] and ground-truth label *y_i_* ∈ {0, 1}, the objective is

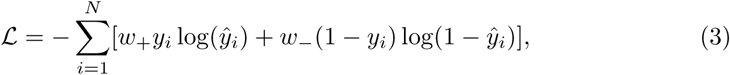

where *w*_+_ and *w*_−_ denote class-specific weights. All parameters of the feature encoder, attention pooling module, and classifier are optimised jointly under this objective.

#### Large-scale patch processing at 20**×** magnification

Whole-slide images processed at 20× magnification typically yield thousands of patches per slide. Forwarding all patches simultaneously through the encoder can lead to memory exhaustion. To maintain stability, we decouple patch-level feature extraction from slide-level aggregation within each forward pass. Patch embeddings are computed in parallel using multi-threaded data loading and streamed into a temporary feature buffer. Once all embeddings for a slide are available, they are assembled into a single bag and passed to the attention pooling and classifier modules.

#### Model parallelism for scalable end-to-end training

The primary memory bottleneck arises during back-propagation. Because gradients must propagate from the classifier through the pooling layer to every patch embedding, activations for all patches must be retained in the computational graph. For slides containing thousands of instances, this can exceed the memory capacity of a single GPU. To enable stable end-to-end optimisation, we employ model parallelism, distributing components of the computational graph across multiple GPUs. Patch-level encoding, attention pooling, and classification are assigned to separate devices, allowing intermediate activations and gradients to be stored across multiple GPU memories. During back-propagation, gradients are computed locally and used to update the corresponding parameter subsets. This strategy permits joint optimisation of the entire framework at high magnification without freezing the encoder or truncating gradients.

#### Training configuration and computational environment

Models were implemented in PyTorch and trained using the Adam optimiser with a weight decay of 5×10^−4^ and an initial learning rate of 1×10^−4^. Training was performed for 100 epochs under three-fold cross-validation, with fold assignments defined at the patient level. Whole-slide image processing was conducted using OpenSlide. Data management and annotation handling were implemented with pandas and NumPy. Statistical analyses were performed using SciPy. Visualisations were generated with Seaborn. All experiments were executed on a multi-GPU system comprising four NVIDIA A100 GPUs (80 GB VRAM each) and 512 GB of system memory.

### Detailed Survival Analysis

Overall survival data were obtained from the TCGA Pan-Cancer Clinical Dataset [31] through NCI Genomic Data Commons [32]. Survival data were available for 1040 out of the 1049 patients used in this study, spanning 12 TCGA cohorts. Prediction scores were binarised to predict ecDNA status using a threshold of 0.5. For patients with multiple WSIs, their maximum AMIE prediction score was used, as we considered a tumour with a single section with ecDNA to be an ecDNA-positive tumour. Kaplan-Meier curves, log-rank tests, and Cox proportional hazard models were performed using Lifelines [33]. Cox models were stratified by tumour type and used ecDNA-negative tumours as the reference category.

### Oncogene-stratified Performance Analysis

In histopathological image analysis, understanding the connection between a model’s predictions and underlying biological factors is essential for interpretability and clinical trust. While our framework predicts ecDNA status from WSIs, a natural curiosity arises: could these predictions also reflect the influence of specific oncogenic drivers? Since ecDNA is often associated with oncogene amplification, we hypothesized that certain oncogenes might, through their presence, leave subtle but detectable morphological signatures that the model could implicitly learn. If such a connection exists, it would suggest a biologically meaningful correspondence between genomic alterations and their manifestation in tissue architecture.

To test this hypothesis, we investigated whether the model’s performance in predicting ecDNA varied with the amplification status of frequently occurring oncogenes across cancer types. This raised the question of whether the presence of a particular oncogene influences the model’s ability to classify ecDNA status from histology. We framed this as a hypothesis test: *the null assumes no performance difference across oncogene-stratified groups, while the alternative suggests that certain amplifications modulate morphology in ways that affect prediction accuracy*. Building on this, we constructed an oncogene–sample indicator matrix. For each sample *W_i_*, the presence of a candidate oncogene *O_k_* within its associated list of amplified genes is recorded. If *O_k_* is present, the corresponding matrix entry is assigned a value of 1; if other oncogenes are listed but not *O_k_*, the entry is assigned a value of 0. Samples with non-oncogenic focal amplifications are represented by 0, similar to samples with no focal amplifications. For each oncogene *O_k_*, we formed two groups: *O*^+^_*k*_ (samples in which *O_k_* is present) and *O*^−^_*k*_ (samples without *O_k_* but possibly with other oncogenes or unknowns). For example, in the BLCA cohort, 17 of the 86 samples in the amplification cluster had CTTN amplified, forming the *CTTN* ^+^ group, while the remaining 69 made up the *CTTN* ^−^ group. For both subgroups, we computed prediction accuracy and confusion matrices (Supplementary Figure A.3). To assess whether these observed variations in accuracy are statistically meaningful, we used the Mann–Whitney U test. This non-parametric test, well-suited for small or uneven sample sizes, produced a *p*-value of 0.0762, indicating no significant difference in classification performance. This suggests that, despite differing molecular contexts, the model’s ability to identify ecDNA was not influenced in any meaningful way by the presence or absence of CTTN amplification.

To complement the accuracy-based evaluation, we further assessed class-specific performance using sensitivity and specificity metrics, analysed via Fisher’s Exact test. In contrast to the Mann–Whitney U test, which compares distributions of prediction correctness, Fisher’s Exact test evaluates categorical associations in contingency tables and is particularly well-suited for small sample sizes. Applied to the CTTN-stratified subgroups, the test yielded *p*-values of 0.1228 for sensitivity and 0.5303 for specificity, indicating no statistically significant differences. These results reinforce the notion that the model’s performance remained broadly stable across CTTN amplification status. We extended this analysis to the candidate oncogenes in the BLCA cohort. The detailed gene-wise performance metrics alongside their associated p-values from both Mann–Whitney U and Fisher’s exact tests are summarised in Supplementary Figure A.4.

The same procedure was extended to all remaining cancer types (Supplementary Figures A.4–A.15) to determine whether gene-conditioned effects were reproducible across tissues. In addition, a pooled pan-cancer analysis was conducted by aggregating samples across tumour types while preserving oncogene stratification (Supplementary Table A.16).

### Data Availability

All histopathology and genomic amplification data used in this study are publicly available. Digitised H&E WSIs were obtained from TCGA through the GDC Data Portal (https://portal.gdc.cancer.gov). Corresponding focal DNA amplification annotations were retrieved from the AmpliconRepository (https://ampliconrepository.org), where amplification architectures were inferred from WGS data using the AmpliconArchitect framework. This study integrates these publicly accessible resources to construct a refined multi-cancer cohort comprising 1,323 whole-slide images from 1,049 patients across 12 cancer types. No new sequencing or imaging data were generated. Access to TCGA data may require compliance with the GDC data use policies.

### Code Availability

The code used for data preprocessing, slide-level augmentation, model training, evaluation, and statistical analyses is publicly available at https://github.com/manwaarkhd/ amie. The repository contains the implementation of the AMIE framework, scripts to reproduce the primary experiments, and documentation to facilitate reuse and independent validation. The code is provided to support transparency, reproducibility, and further methodological development by the research community.

---

**Table A.1:**
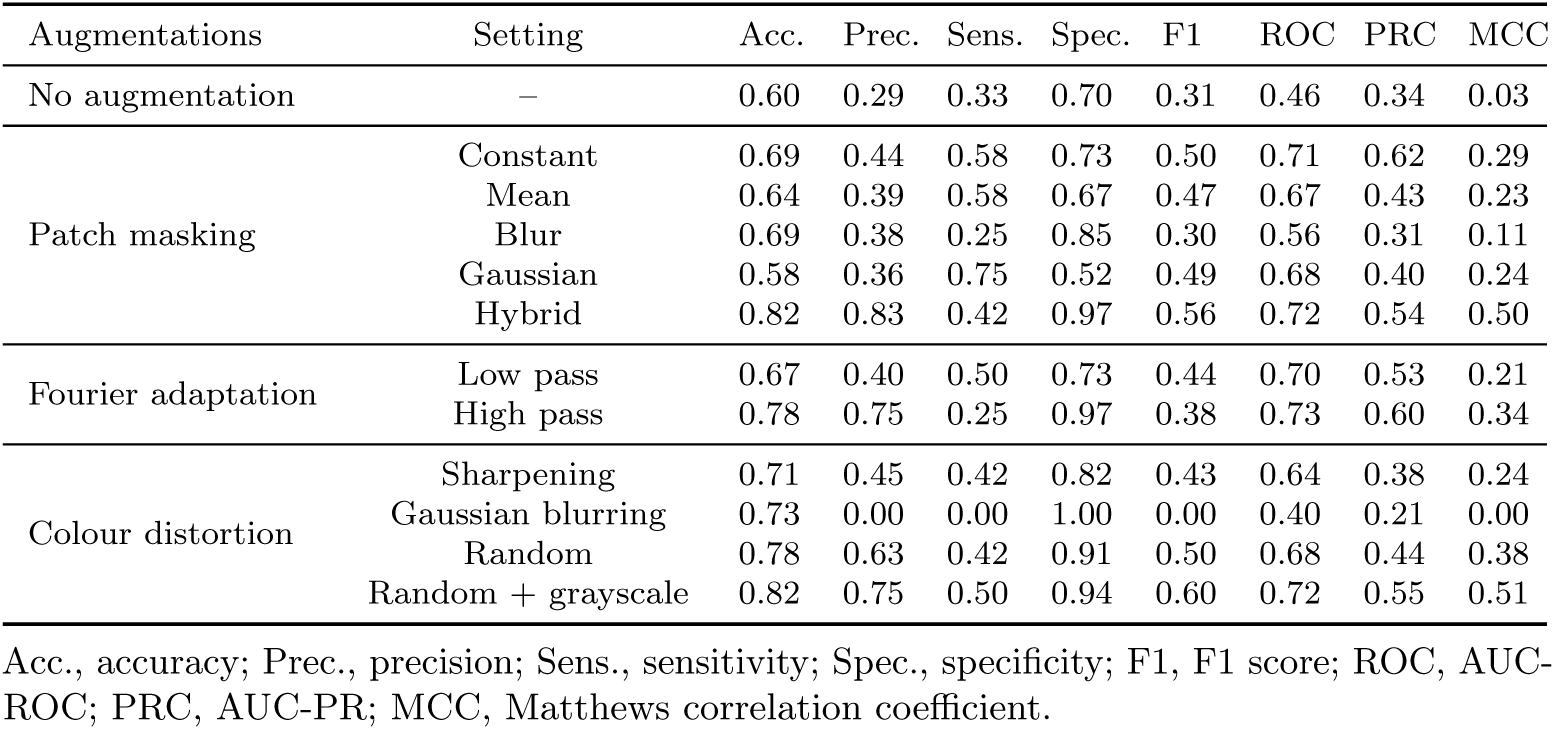
Impact of slide-level augmentation parametrizations on ecDNA-driven tumour classification in TCGA-BLCA.

**Fig. A.1:**
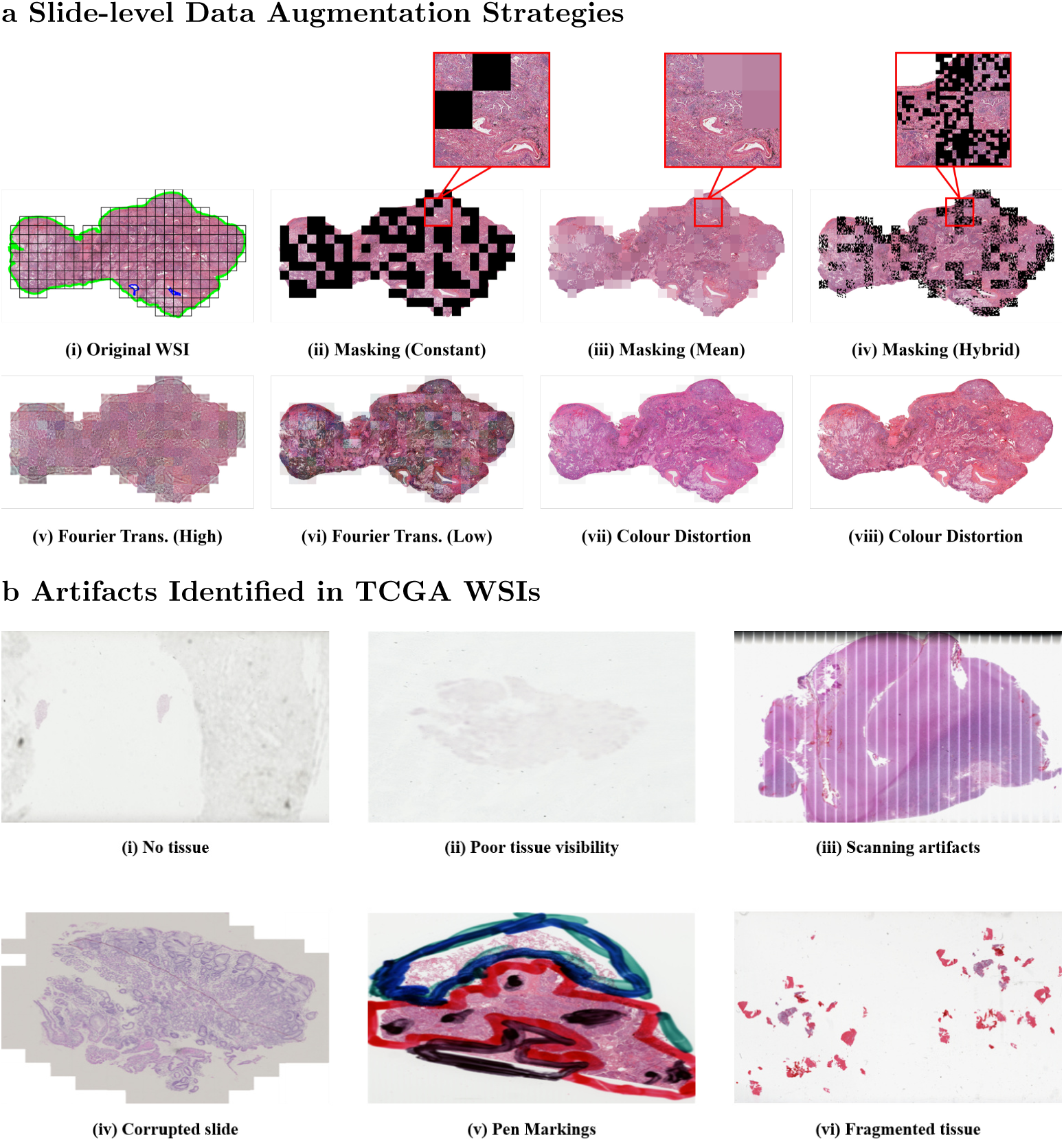
Slide-level quality control and augmentation strategies for robust model training. (a) Overview of coordinated slide-level augmentation strategies, including stochastic patch masking, Fourier-domain perturbations, and stain-aware colour transformations. (b) Representative WSIs illustrating common artifacts identified during quality control, including absent or poorly visible tissue, scanning artifacts, corrupted files, pen markings, and fragmented tissue distribution.

**Fig. A.2:**
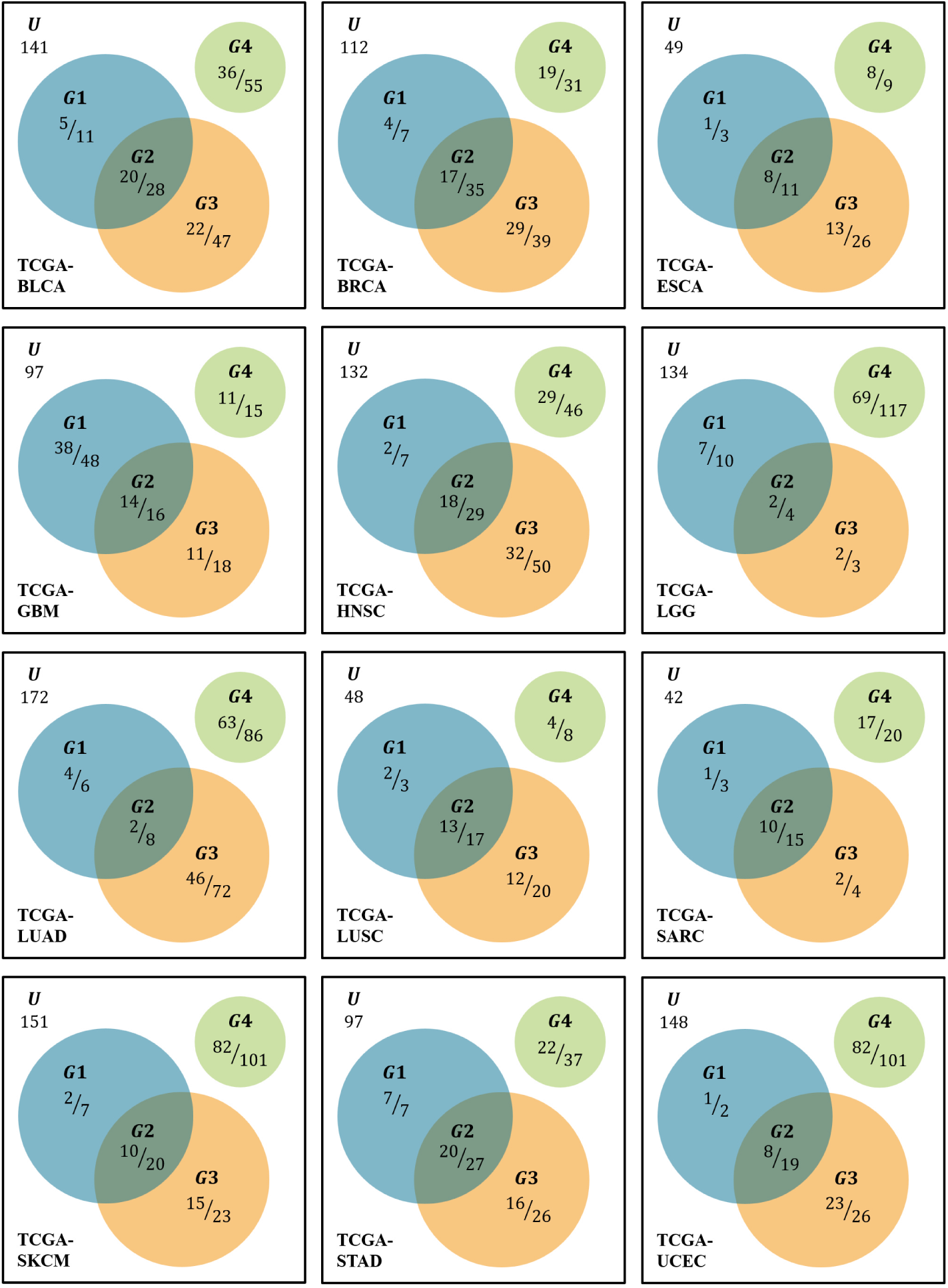
Per-cancer ecDNA classification outcomes stratified by amplification context. For each tumour type, Venn diagrams partition patients into ecDNA-only amplification *G*1, ecDNA co-occurring with other focal amplification architectures *G*2, focal chromosomal amplification without ecDNA *G*3, and no focal amplification *G*4. ecDNA-positive cases correspond to *G*1 ∪ *G*2. Values indicate correctly classified cases relative to total cases within each category. The diagrams illustrate context-dependent variation in classification behaviour across amplification contexts and tumour types.

**Fig. A.3:**
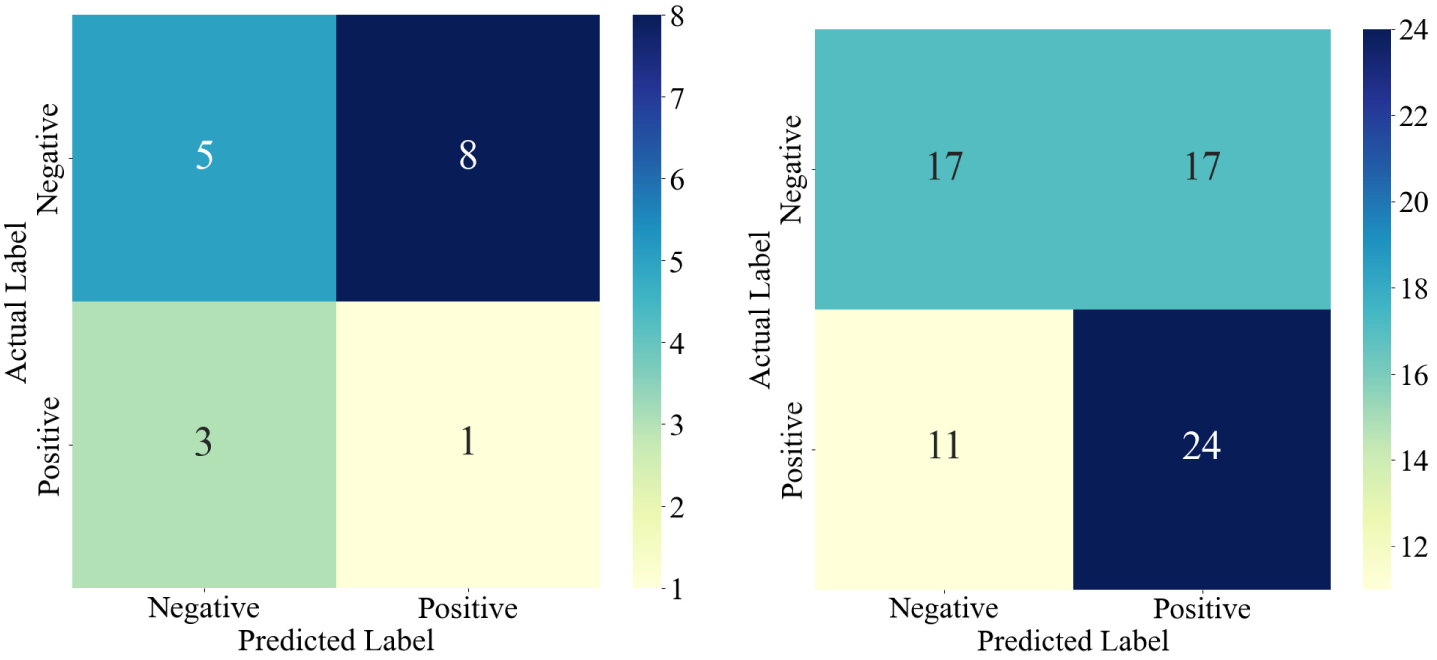
Confusion matrices showing model prediction performance for BLCA samples with and without CTTN amplification. (a) Prediction outcomes for samples with *CTTN* amplification (*CTTN* ^+^). (b) Prediction outcomes for samples without *CTTN* amplification (*CTTN* ^−^).

**Fig. A.4:**
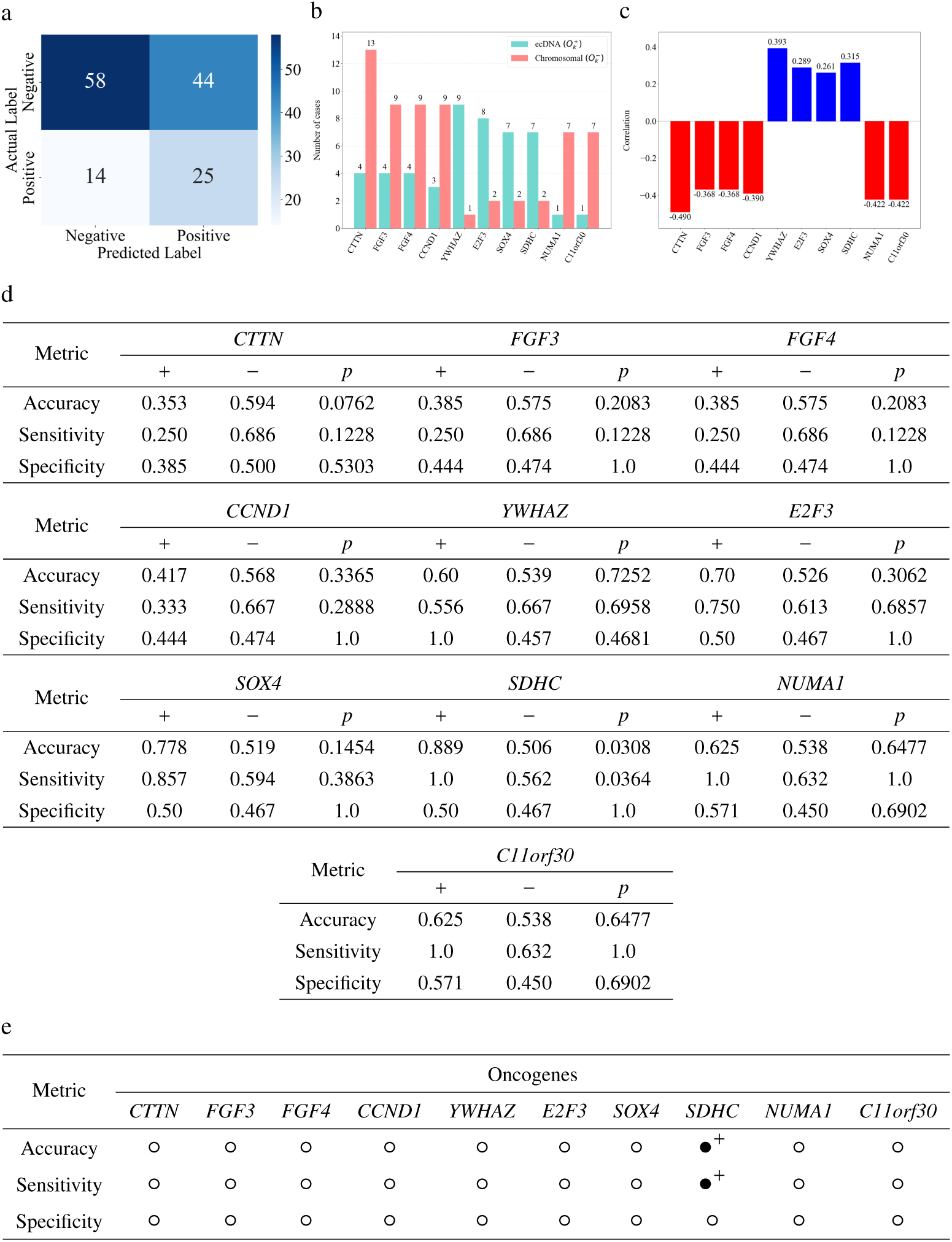
Oncogene-stratified evaluation of ecDNA prediction performance in BLCA. (a) Confusion matrix. (b) ecDNA-stratified oncogenes distribution. (c) Correlation of oncogenes with ecDNA. (d) Oncogene-specific impact on prediction performance in TCGA-BLCA. (e) Summary of statistical significance and directionality across oncogenes.

**Fig. A.5:**
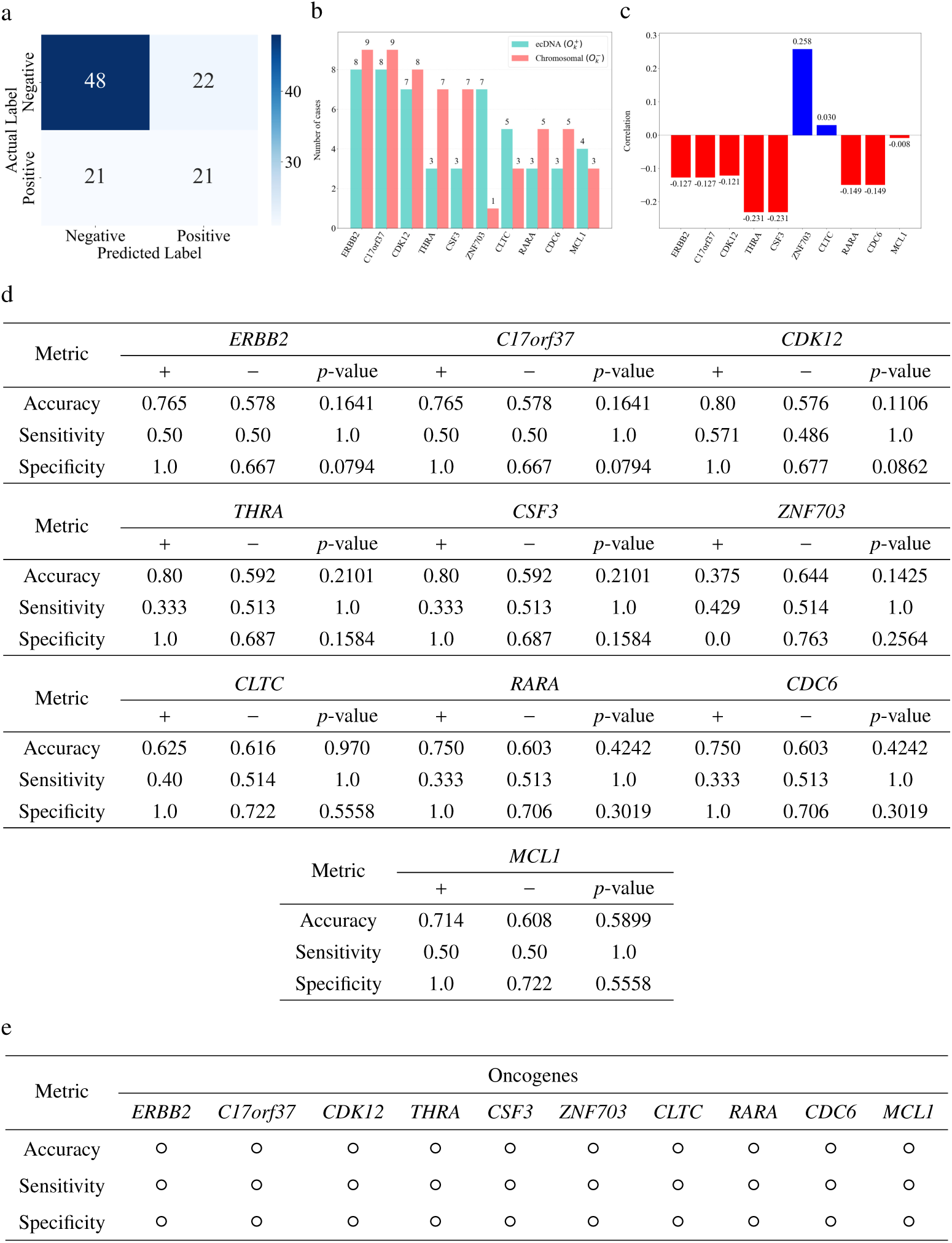
Oncogene-stratified evaluation of ecDNA prediction performance in BRCA. (a) Confusion matrix. (b) ecDNA-stratified oncogenes distribution. (c) Correlation of oncogenes with ecDNA. (d) Oncogene-specific impact on prediction performance in TCGA-BRCA. (e) Summary of statistical significance and directionality across oncogenes.

**Fig. A.6:**
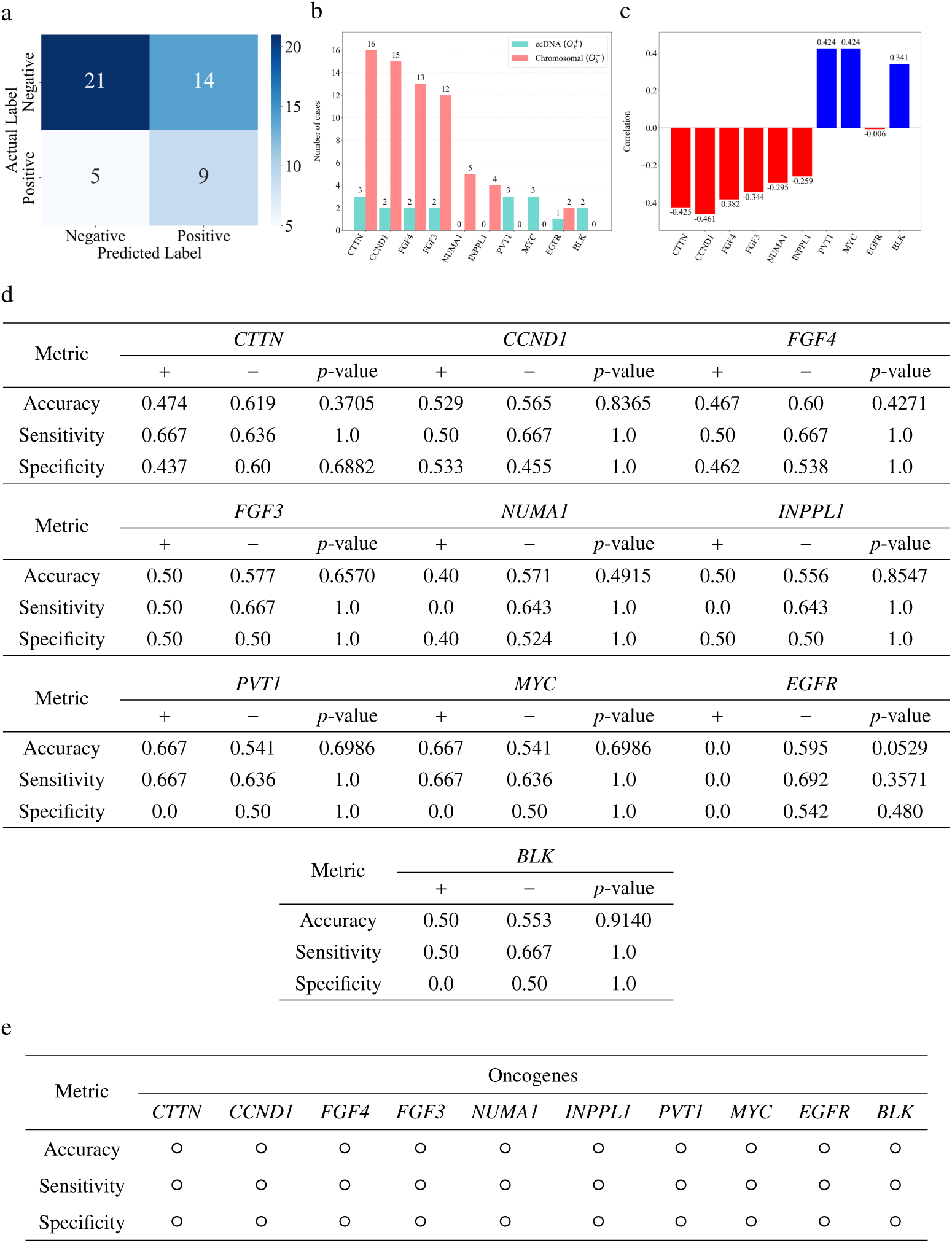
Oncogene-stratified evaluation of ecDNA prediction performance in ESCA. (a) Confusion matrix. (b) ecDNA-stratified oncogenes distribution. (c) Correlation of oncogenes with ecDNA. (d) Oncogene-specific impact on prediction performance in TCGA-ESCA. (e) Summary of statistical significance and directionality across oncogenes.

**Fig. A.7:**
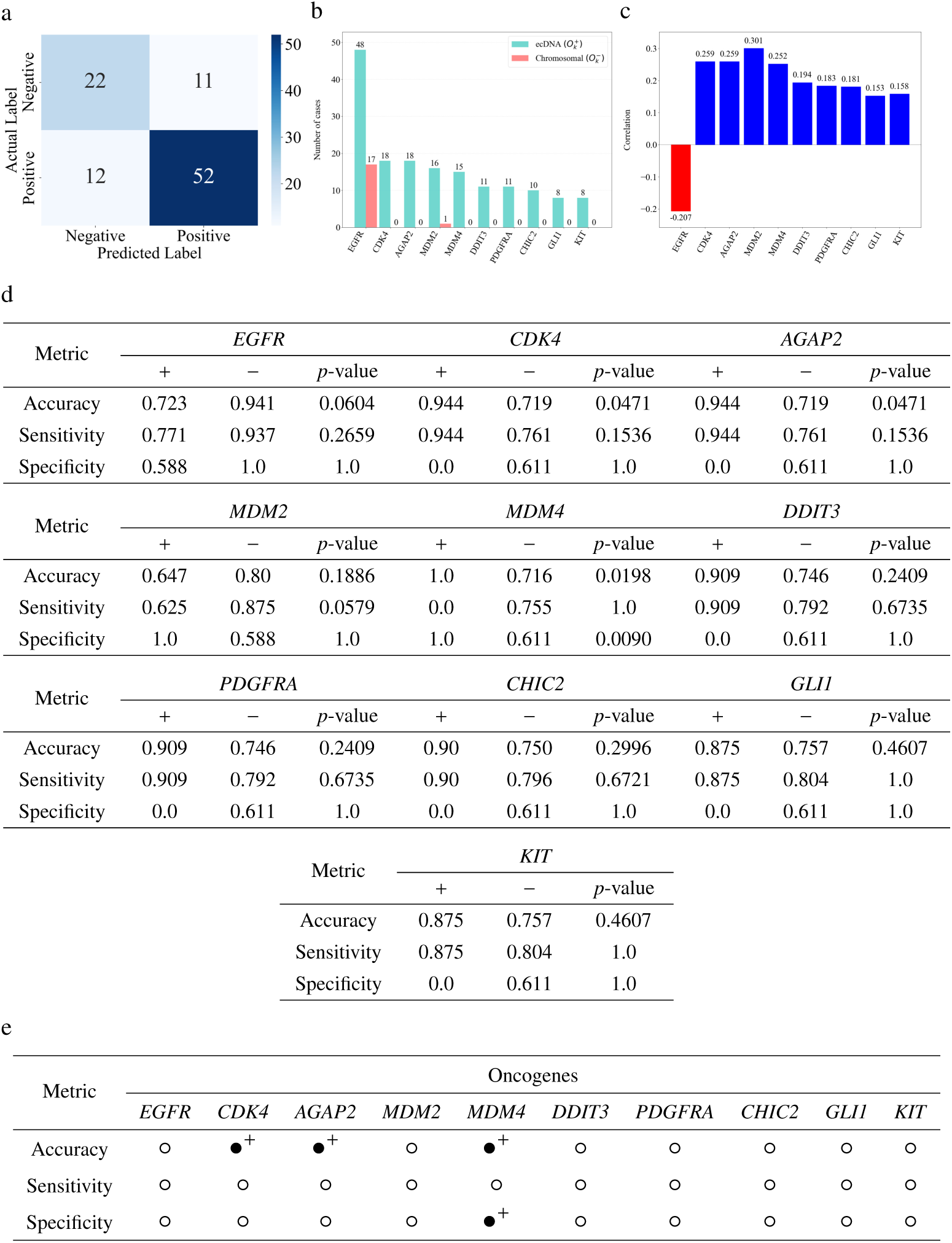
Oncogene-stratified evaluation of ecDNA prediction performance in GBM. (a) Confusion matrix. (b) ecDNA-stratified oncogenes distribution. (c) Correlation of oncogenes with ecDNA. (d) Oncogene-specific impact on prediction performance in TCGA-GBM. (e) Summary of statistical significance and directionality across oncogenes.

**Fig. A.8:**
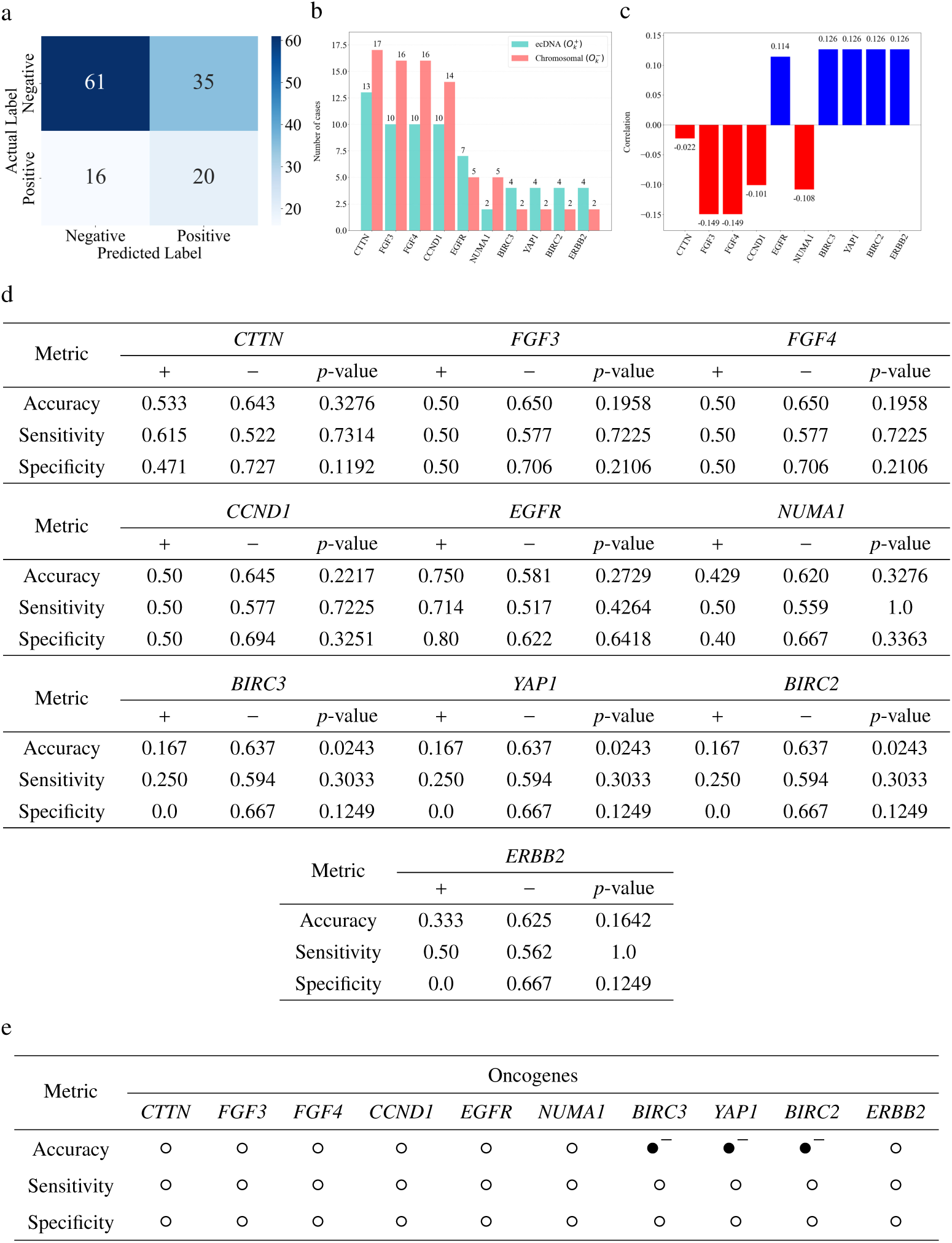
Oncogene-stratified evaluation of ecDNA prediction performance in HNSC. (a) Confusion matrix. (b) ecDNA-stratified oncogenes distribution. (c) Correlation of oncogenes with ecDNA. (d) Oncogene-specific impact on prediction performance in TCGA-HNSC. (e) Summary of statistical significance and directionality across oncogenes.

**Fig. A.9:**
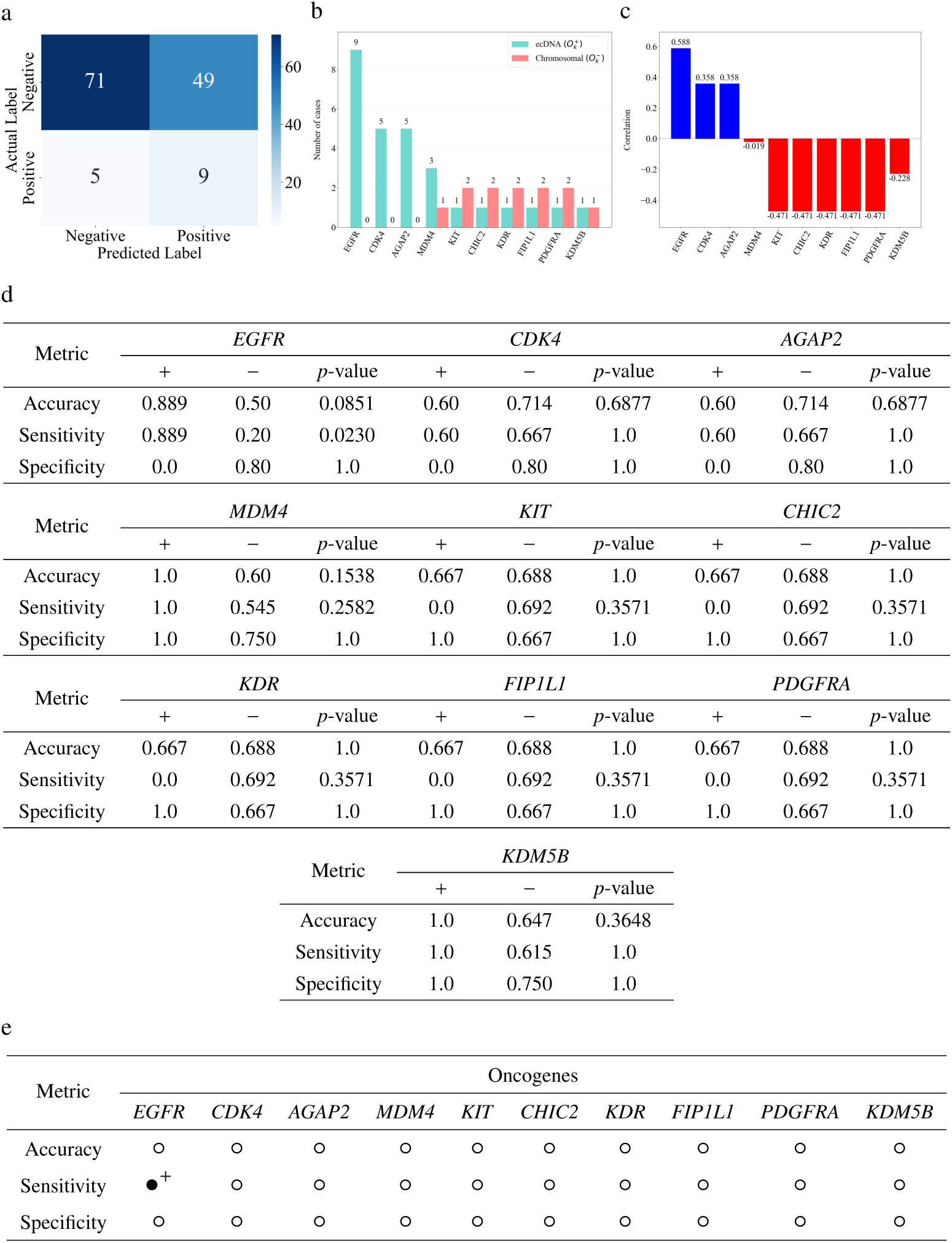
Oncogene-stratified evaluation of ecDNA prediction performance in LGG. (a) Confusion matrix. (b) ecDNA-stratified oncogenes distribution. (c) Correlation of oncogenes with ecDNA. (d) Oncogene-specific impact on prediction performance in TCGA-LGG. (e) Summary of statistical significance and directionality across oncogenes.

**Fig. A.10:**
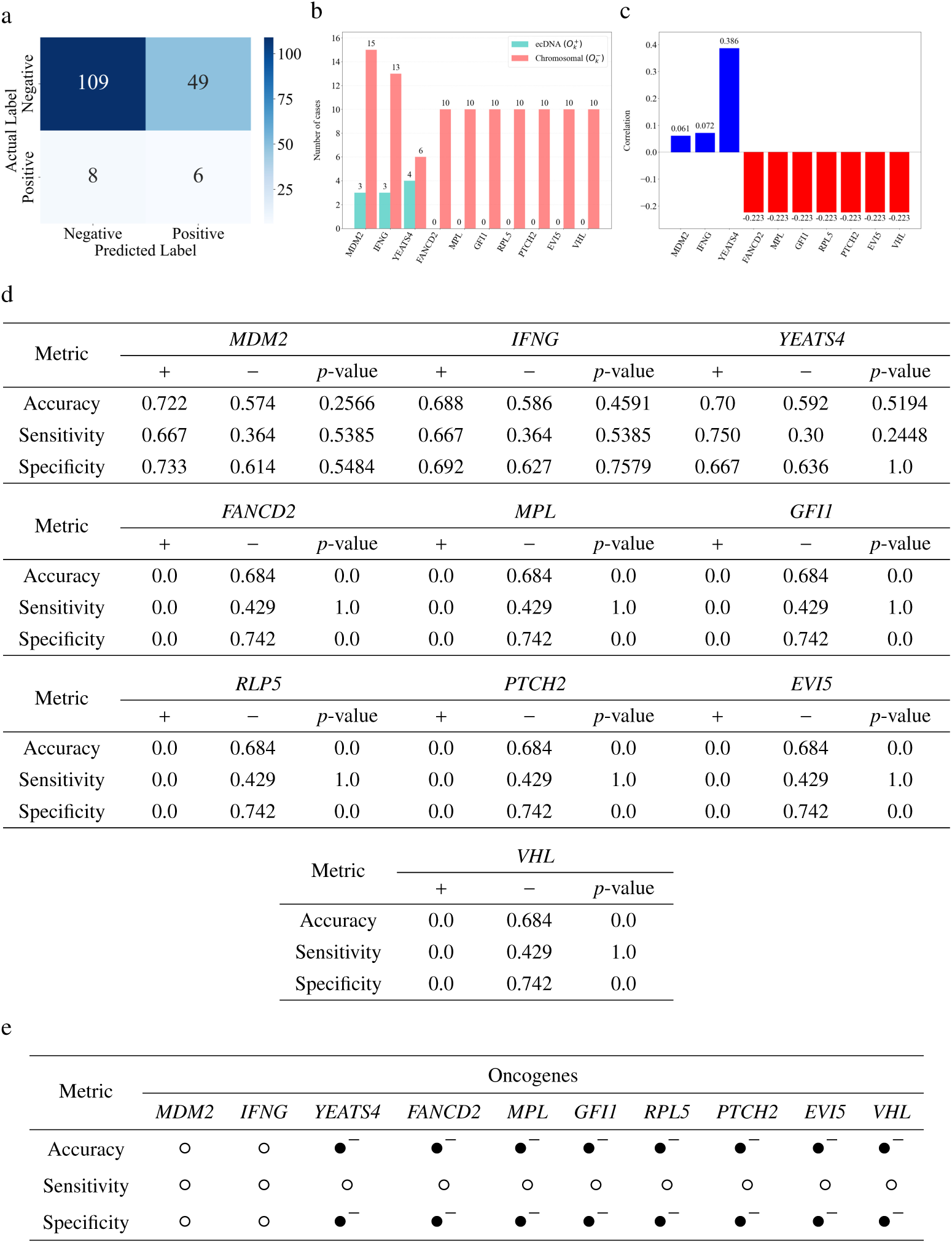
Oncogene-stratified evaluation of ecDNA prediction performance in LUAD. (a) Confusion matrix. (b) ecDNA-stratified oncogenes distribution. (c) Correlation of oncogenes with ecDNA. (d) Oncogene-specific impact on prediction performance in TCGA-LUAD. (e) Summary of statistical significance and directionality across oncogenes.

**Fig. A.11:**
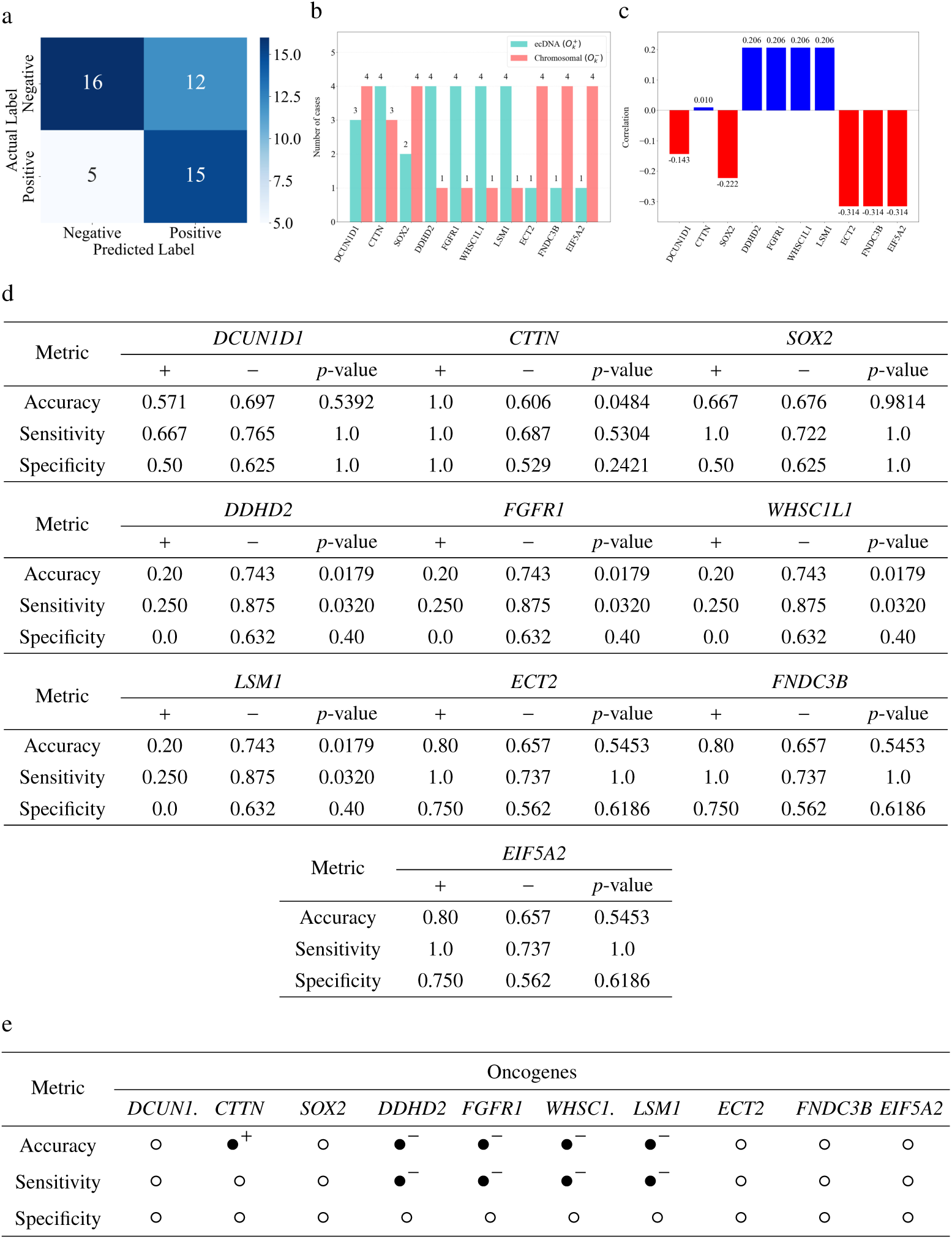
Oncogene-stratified evaluation of ecDNA prediction performance in LUSC. (a) Confusion matrix. (b) ecDNA-stratified oncogenes distribution. (c) Correlation of oncogenes with ecDNA. (d) Oncogene-specific impact on prediction performance in TCGA-LUSC. (e) Summary of statistical significance and directionality across oncogenes. For visual clarity, *DCUN1D1* and *WHSC1L1* are displayed as *DCUN1* and *WHSC1*, respectively. _38_

**Fig. A.12:**
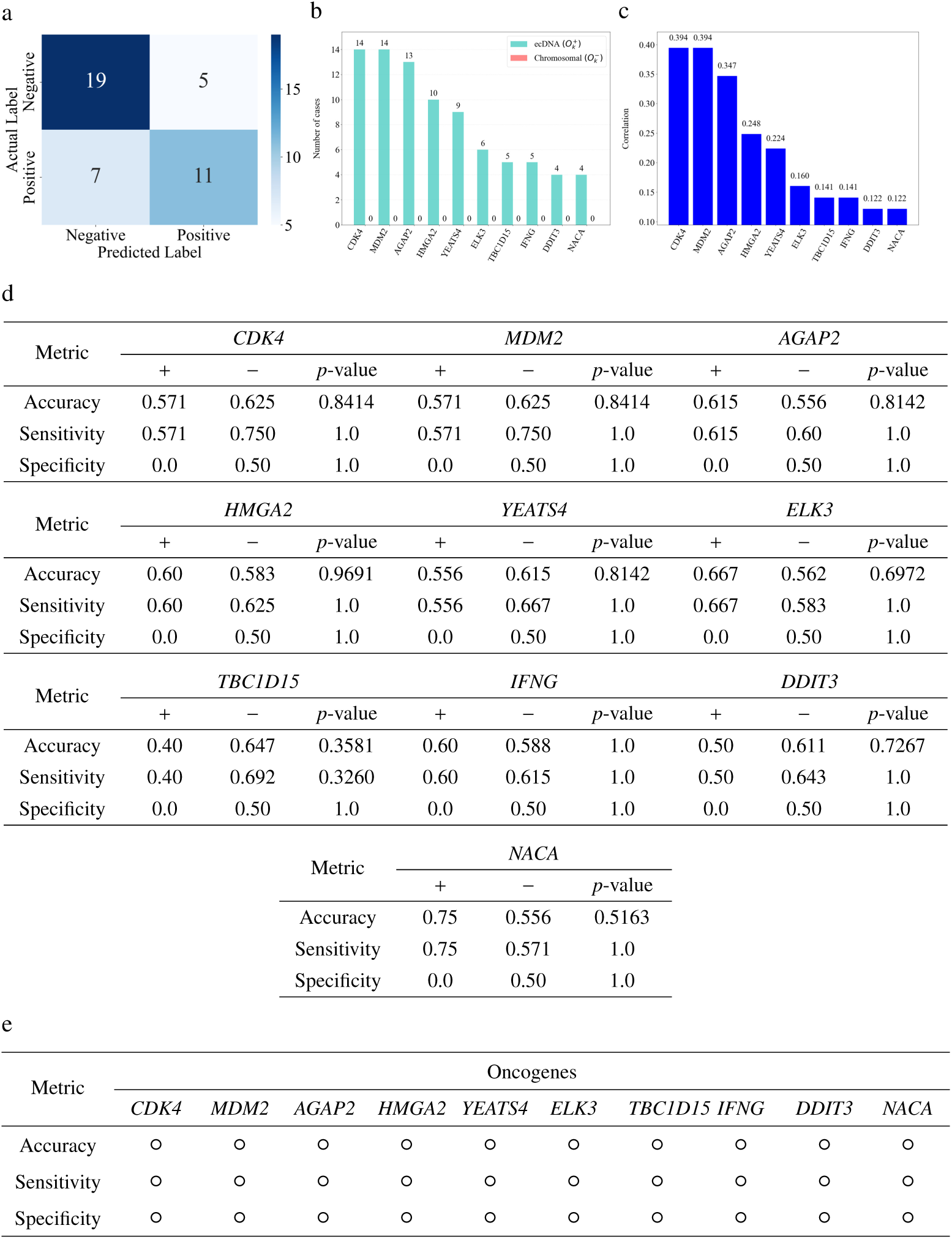
Oncogene-stratified evaluation of ecDNA prediction performance in SARC. (a) Confusion matrix. (b) ecDNA-stratified oncogenes distribution. (c) Correlation of oncogenes with ecDNA. (d) Oncogene-specific impact on prediction performance in TCGA-SARC. (e) Summary of statistical significance and directionality across oncogenes.

**Fig. A.13:**
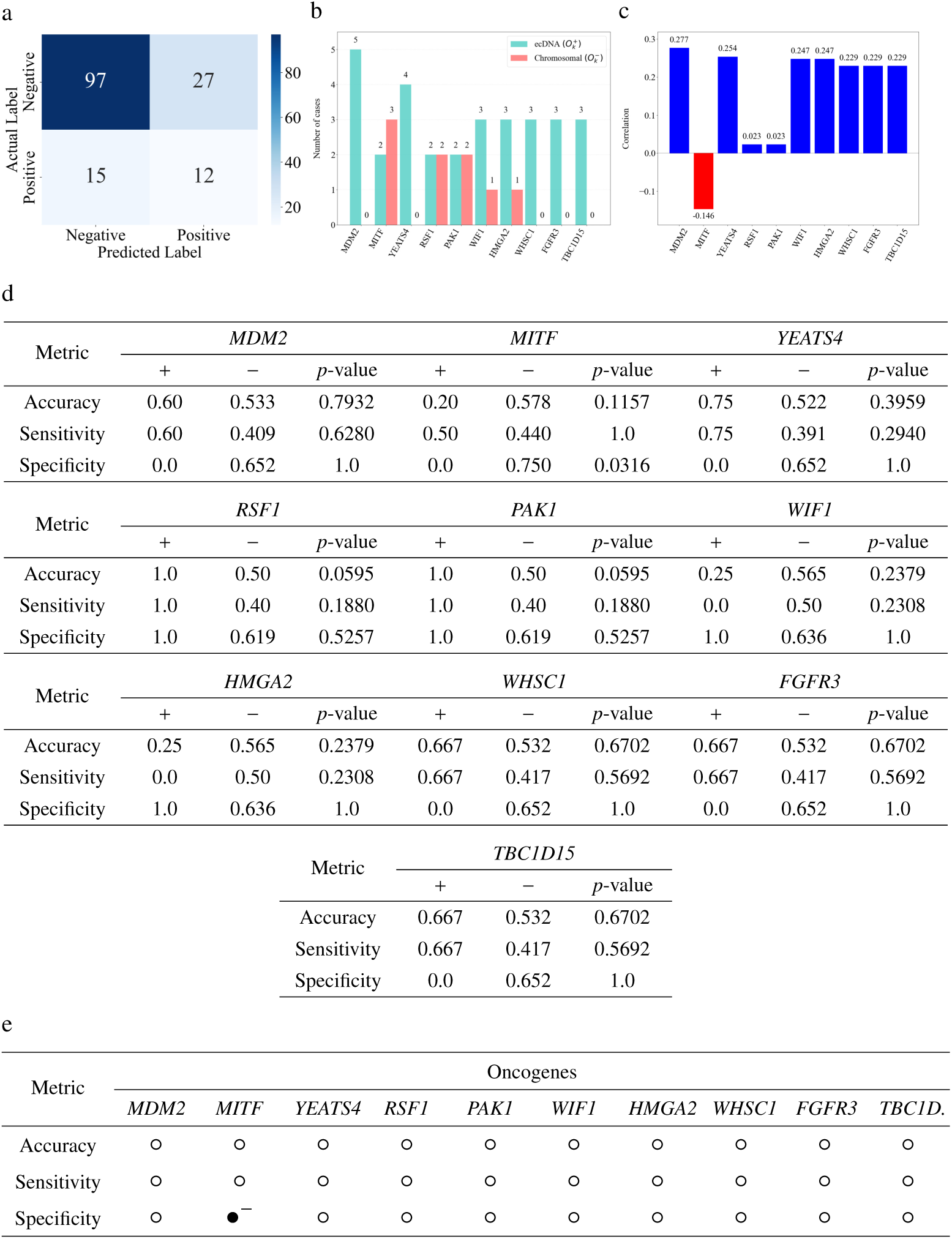
Oncogene-stratified evaluation of ecDNA prediction performance in SKCM. (a) Confusion matrix. (b) ecDNA-stratified oncogenes distribution. (c) Correlation of oncogenes with ecDNA. (d) Oncogene-specific impact on prediction performance in TCGA-SKCM. (e) Summary of statistical significance and directionality across oncogenes. For visual clarity, *TBC1D15* is displayed as *TBC1D*.

**Fig. A.14:**
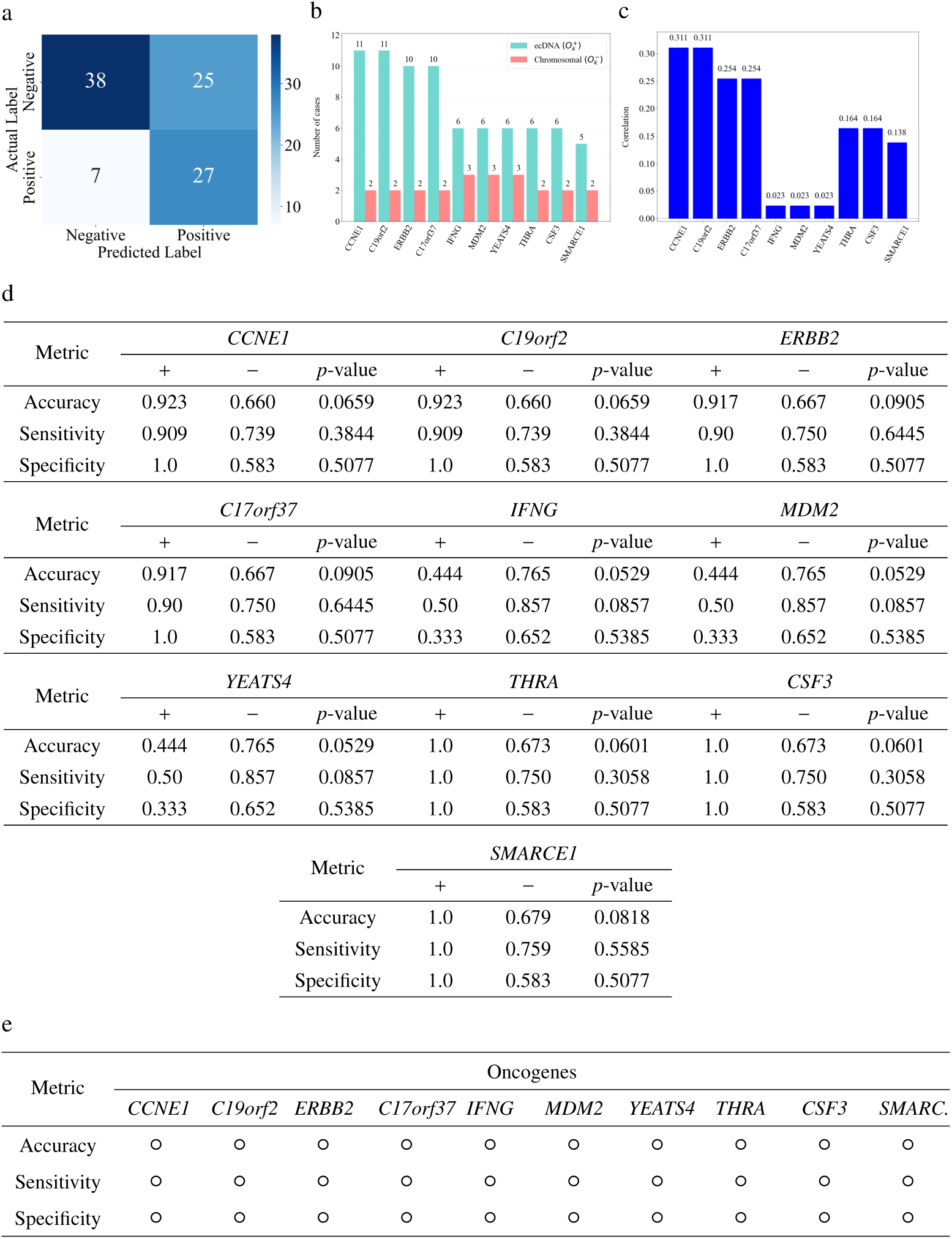
Oncogene-stratified evaluation of ecDNA prediction performance in STAD. (a) Confusion matrix. (b) ecDNA-stratified oncogenes distribution. (c) Correlation of oncogenes with ecDNA. (d) Oncogene-specific impact on prediction performance in TCGA-STAD. (e) Summary of statistical significance and directionality across oncogenes. For visual clarity, *SMARCE1* is displayed as *SMARC*.

**Fig. A.15:**
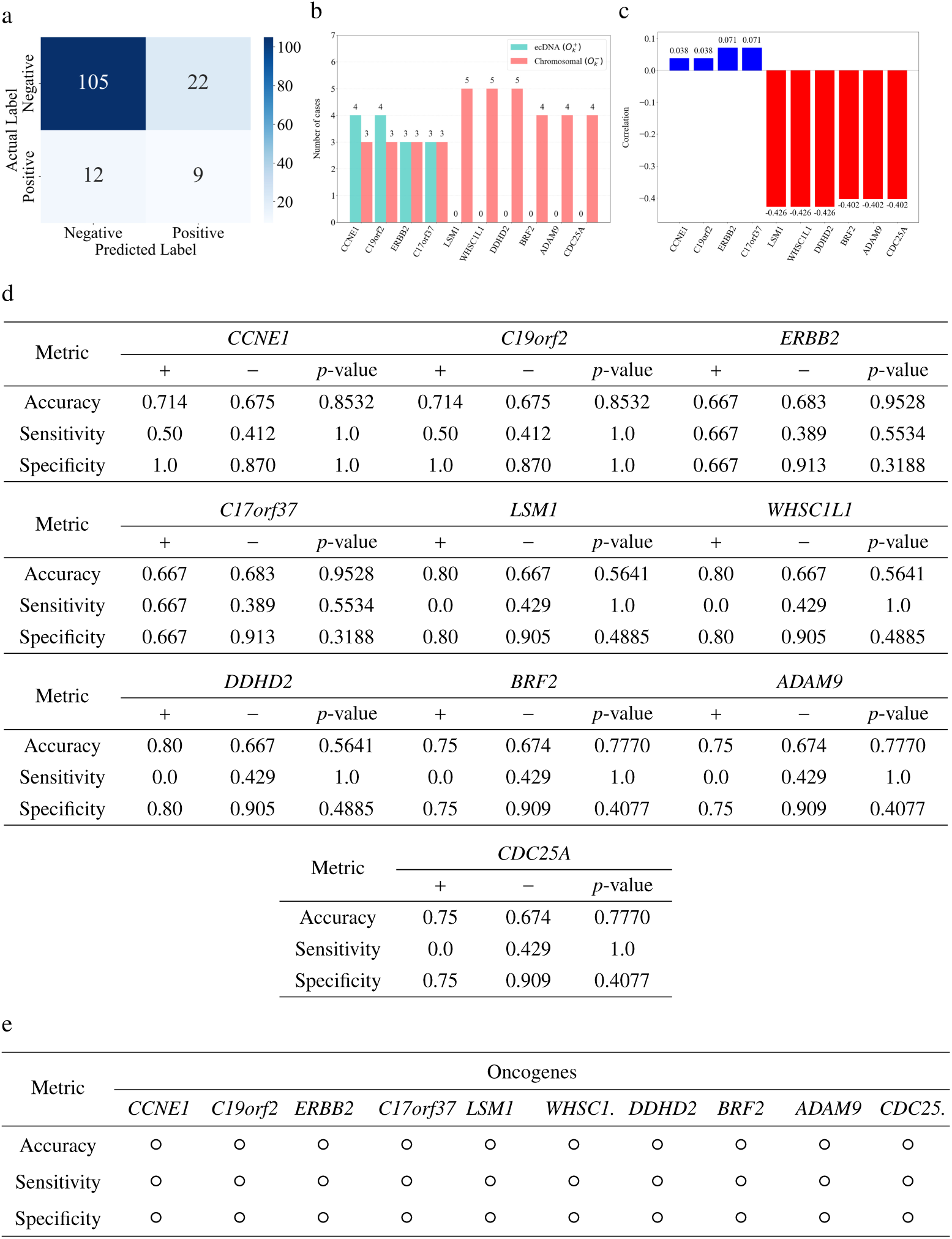
Oncogene-stratified evaluation of ecDNA prediction performance in UCEC. (a) Confusion matrix. (b) ecDNA-stratified oncogenes distribution. (c) Correlation of oncogenes with ecDNA. (d) Oncogene-specific impact on prediction performance in TCGA-UCEC. (e) Summary of statistical significance and directionality across oncogenes. For visual clarity, gene labels are abbreviated as *WHSC1* (*WHSC1L1*) and *CDC25* (*CDC25A*).

**Fig. A.16:**
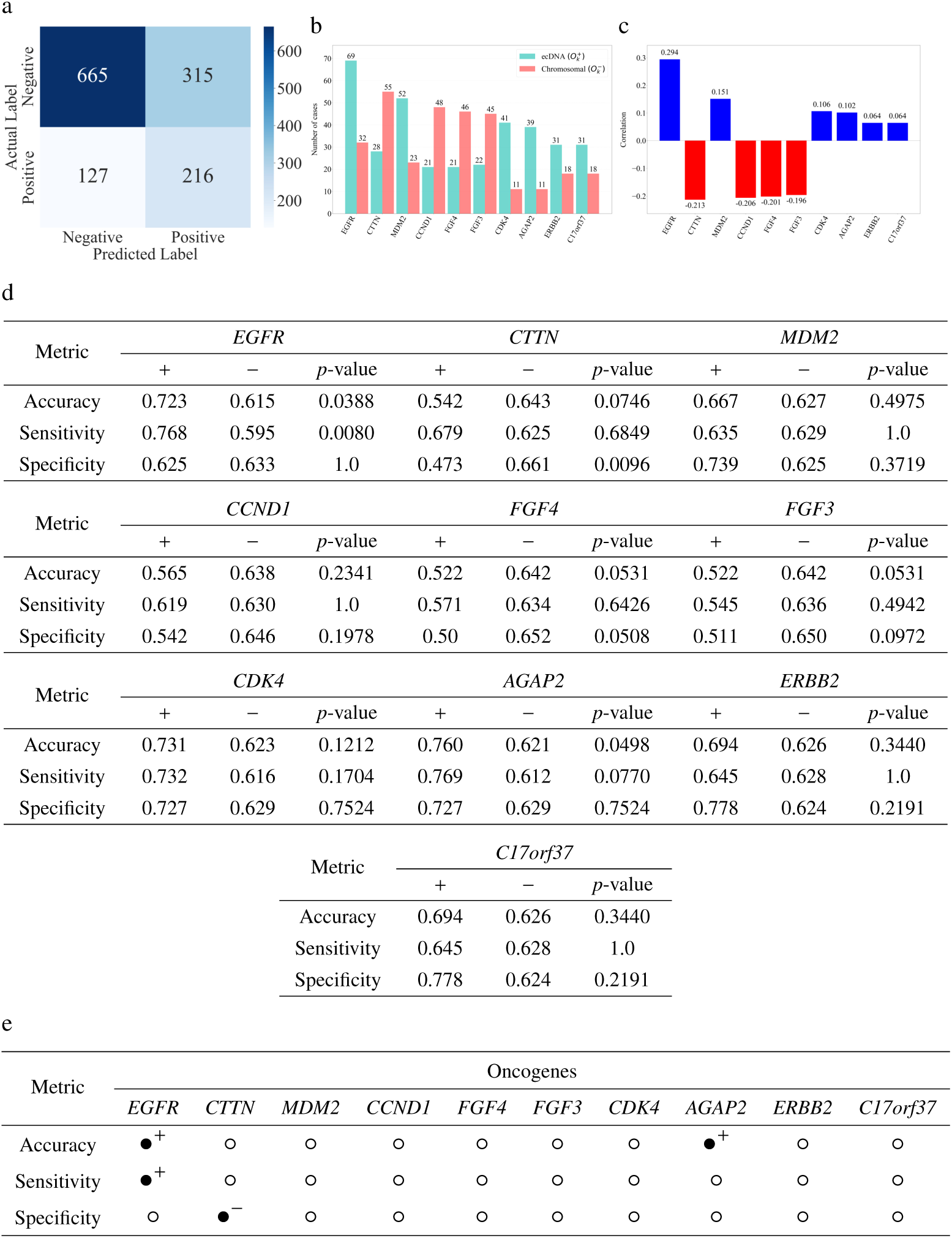
Oncogene-stratified evaluation of ecDNA prediction performance across TCGA. (a) Confusion matrix. (b) ecDNA-stratified oncogenes distribution. (c) Correlation of oncogenes with ecDNA. (d) Oncogene-specific impact on prediction performance across TCGA. (e) Summary of statistical significance and directionality across oncogenes.

## Notes

### Competing Interest Statement

The authors have declared no competing interest.

## References

[1] Turner, K. M. et al. Extrachromosomal oncogene amplification drives tumour evolution and genetic heterogeneity. Nature 543, 122–125 (2017).

[2] Wu, S. et al. Circular ecdna promotes accessible chromatin and high oncogene expression. Nature 575, 699–703 (2019).

[3] Verhaak, R. G., Bafna, V. & Mischel, P. S. Extrachromosomal oncogene amplification in tumour pathogenesis and evolution. Nature Reviews Cancer 19, 283–288 (2019).

[4] Yan, X., Mischel, P. & Chang, H. Extrachromosomal dna in cancer. Nature Reviews Cancer 24, 261–273 (2024).

[5] Kim, H. et al. Extrachromosomal dna is associated with oncogene amplification and poor outcome across multiple cancers. Nature genetics 52, 891–897 (2020).

[6] Bailey, C. et al. Origins and impact of extrachromosomal dna. Nature 635, 193–200 (2024).

[7] Chapman, O. S. et al. Circular extrachromosomal dna promotes tumor heterogeneity in high-risk medulloblastoma. Nature genetics 55, 2189–2199 (2023).

[8] Chapman, O. S., et al. Extrachromosomal dna associates with poor survival across a broad spectrum of childhood solid tumors. *MedRxiv* (2025).

[9] Koche, R. P. et al. Extrachromosomal circular dna drives oncogenic genome remodeling in neuroblastoma. Nature genetics 52, 29–34 (2020).

[10] Luebeck, J. et al. Extrachromosomal dna in the cancerous transformation of barrett’s oesophagus. Nature 616, 798–805 (2023).

[11] Karami Fath, M., et al. Revisiting characteristics of oncogenic extrachromosomal dna as mobile enhancers on neuroblastoma and glioma cancers. Cancer Cell International 22, 200 (2022).

[12] Deshpande, V. et al. Exploring the landscape of focal amplifications in cancer using ampliconarchitect. Nature communications 10, 392 (2019).

[13] Song, A. H. et al. Artificial intelligence for digital and computational pathology. Nature Reviews Bioengineering 1, 930–949 (2023).

[14] Kather, J. N. et al. Pan-cancer image-based detection of clinically actionable genetic alterations. Nature cancer 1, 789–799 (2020).

[15] Fiorini, E. et al. Myc ecdna promotes intratumour heterogeneity and plasticity in pdac. Nature 1–10 (2025).

[16] Zhao, B. et al. Oncogenic drivers shape the tumor microenvironment in human gliomas. bioRxiv 2025–05 (2025).

[17] Weinstein, J. N. et al. The cancer genome atlas pan-cancer analysis project. Nature genetics 45, 1113–1120 (2013).

[18] Verhaak, R. G. et al. Integrated genomic analysis identifies clinically relevant subtypes of glioblastoma characterized by abnormalities in pdgfra, idh1, egfr, and nf1. Cancer cell 17, 98–110 (2010).

[19] Mazzoleni, A. et al. Chromosomal instability: a key driver in glioma pathogenesis and progression. European Journal of Medical Research 29, 451 (2024).

[20] Hung, K. L. et al. ecdna hubs drive cooperative intermolecular oncogene expression. Nature 600, 731–736 (2021).

[21] Vorontsov, E. et al. A foundation model for clinical-grade computational pathology and rare cancers detection. Nature medicine 30, 2924–2935 (2024).

[22] Chen, R. J. et al. Towards a general-purpose foundation model for computational pathology. Nature Medicine 30, 850–862 (2024).

[23] Wang, X. et al. Transformer-based unsupervised contrastive learning for histopathological image classification. Medical image analysis 81, 102559 (2022).

[24] Graham, S. et al. Hover-net: Simultaneous segmentation and classification of nuclei in multi-tissue histology images. Medical image analysis 58, 101563 (2019).

[25] Kocak, B. et al. Evaluation metrics in medical imaging ai: fundamentals, pitfalls, misapplications, and recommendations. European Journal of Radiology Artificial Intelligence 100030 (2025).

[26] Lu, M. Y. et al. Data-efficient and weakly supervised computational pathology on whole-slide images. Nature biomedical engineering 5, 555–570 (2021).

[27] Otsu, N. et al. A threshold selection method from gray-level histograms. Automatica 11, 23–27 (1975).

[28] Gonzales, R. C. & Wintz, P. Digital image processing (Addison-Wesley Longman Publishing Co., Inc., 1987).

[29] Suzuki, S. et al. Topological structural analysis of digitized binary images by border following. Computer vision, graphics, and image processing 30, 32–46 (1985).

[30] Campanella, G. et al. Clinical-grade computational pathology using weakly supervised deep learning on whole slide images. Nature medicine 25, 1301–1309 (2019).

[31] Liu, J. et al. An integrated tcga pan-cancer clinical data resource to drive high-quality survival outcome analytics. Cell 173, 400–416 (2018).

[32] Heath, A. P. et al. The nci genomic data commons. Nature genetics 53, 257–262 (2021).

[33] Davidson-Pilon, C. lifelines: survival analysis in python. Journal of Open Source Software 4, 1317 (2019).

